# Patient phenotypes and their relation to TNFα signaling and immune cell composition in critical illness and autoimmune disease

**DOI:** 10.1101/2023.10.30.564631

**Authors:** Vinod Krishna, Homayon Banie, Nádia Conceição-Neto, Yoshihiko Murata, Inge Verbrugge, Vladimir Trifonov, Roxana Martinez, Vasumathy Murali, Yu-Chi Lee, Richard D May, Isabel Nájera, Andrew Fowler, Chris Ka Fai Li

## Abstract

**Rationale:** TNFα inhibitors have shown promise in reducing mortality in hospitalized COVID-19 patients; one hypothesis explaining the limited clinical efficacy is patient heterogeneity in the TNFα pathway.

**Methods:** We evaluated the effect of TNFα inhibitors in a mouse model of LPS-induced acute lung injury. Using machine learning we attempted predictive enrichment of TNFα signaling in patients with either ARDS or sepsis. We examined biological factors that drive heterogeneity in host responses to critical infection and their relation to clinical outcomes.

**Results:** In mice, LPS induced TNFα–dependent neutrophilia, alveolar permeability and endothelial injury. In humans, TNFα pathway activation was significantly increased in peripheral blood of patients with critical illnesses and associated with the presence of mature neutrophils across critical illnesses and several autoimmune conditions. Machine learning using a gene signature separated patients into 5 phenotypes; one was a hyper-inflammatory, interferon-associated phenotype enriched for increased TNFα pathway activation and conserved across critical illnesses and autoimmune diseases. Cell subset profiles segregated severely ill patients into neutrophil-subset-dependent groups that were enriched for disease severity, demonstrating the importance of neutrophils in the immune response in critical illness.

**Conclusions:** TNFα signaling and mature neutrophils are associated with a hyper-inflammatory phenotype of patients, shared across critical illness and autoimmune disease. This phenotyping provides a personalized medicine hypothesis to test anti-TNFα therapy in severe respiratory illness.

**Graphical abstract:** 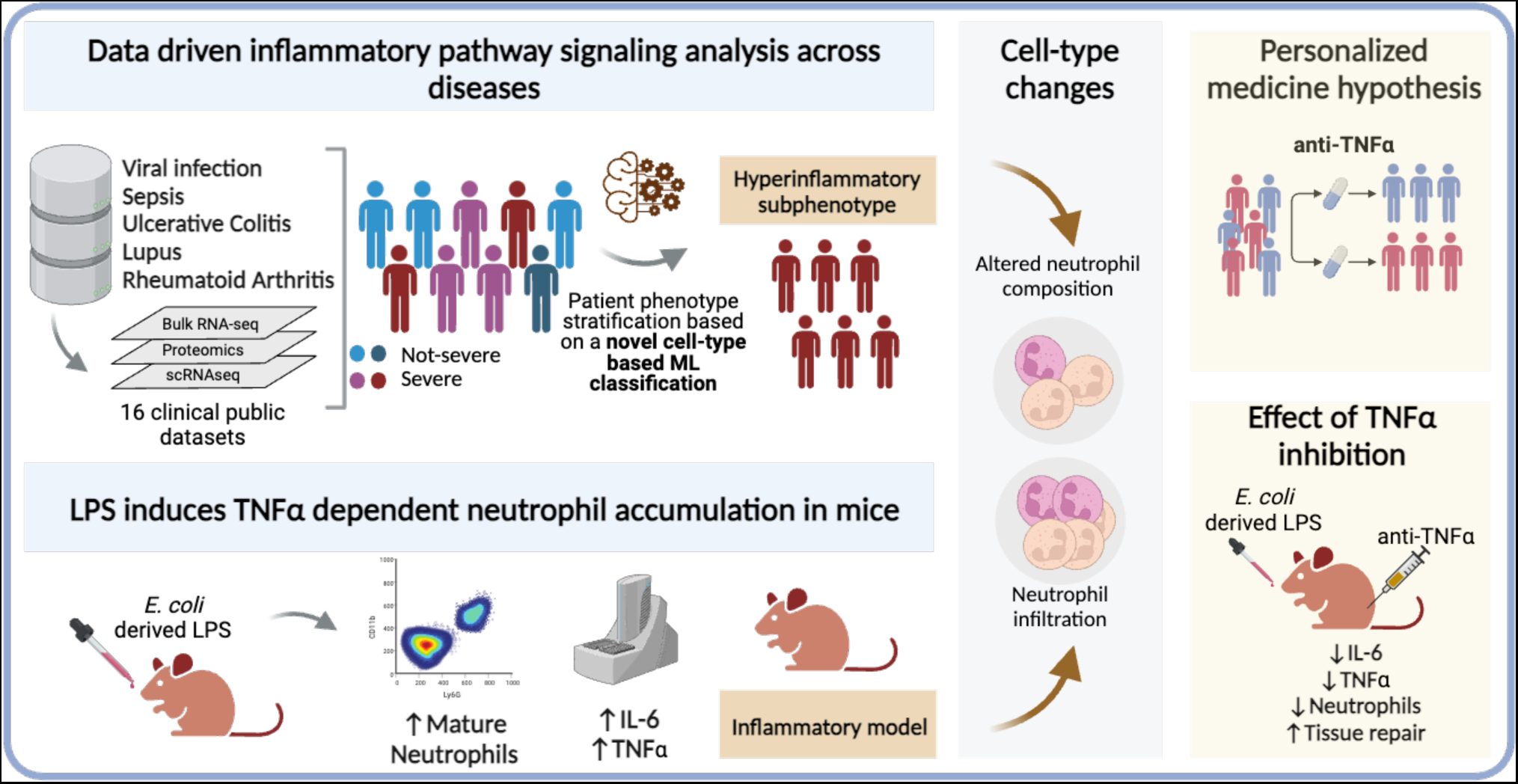

## Introduction

Acute respiratory distress syndrome (ARDS) and sepsis are heterogeneous critical illnesses that have multiple causative factors, with infection being the most common [1]. Therapies for these syndromes have not achieved significant clinical benefit [2, 3]. The considerable heterogeneity in these critical illnesses is believed to be a major reason for this failure, and there is interest in precision medicine approaches to treat both sepsis and ARDS [1–3].

Retrospective analysis of data from clinical trials identified two ARDS phenotypes, a hyperinflammatory phenotype associated with elevated levels of plasma inflammatory proteins (interleukin (IL)-6, soluble TNFR1 and IL-8) and enhanced mortality relative to the other, hypoinflammatory phenotype [4–7]. A clinical trial of ARDS patients on lung-protective mechanical ventilation showed, relative to controls, a reduction in bronchoalveolar lavage fluid (BALF) neutrophils, and BALF and plasma IL-6 and TNFα levels [8]. Analysis of sepsis clinical studies identified homogenous phenotypes, including phenotypes with high inflammatory profiles [1, 2]. However, clinical development of therapies validated with animal models have been unsuccessful, perhaps due to limited ability of models to mimic heterogenous mechanisms seen in humans [1].

TNFα is a key pro-inflammatory mediator in sepsis [9]. A meta-analysis of various studies assessing patients with severe sepsis (without shock) showed that anti-TNF-α inhibitors could reduce mortality [10]. Severely ill patients due to COVID-19 infection showed increased levels of circulating pro-inflammatory cytokines such as TNFα, IL-6, interferon-γ, and IL-18 [11]. Clinical studies assessing TNFα-inhibition or IL-6R blockade in COVID-19 patients showed reduced mortality [12], with a trial of anti-TNF-α infliximab in patients hospitalized with COVID demonstrating improvement in 14-, 28- and 60-day mortality rates relative to placebo [13].

We aimed to develop drug discovery processes to deliver therapeutics for clinical testing in ARDS, with the TNFα axis as a first candidate. We developed both mechanistic preclinical models and computational analyses to enable precision medicine strategies for clinical testing of drug candidates. We established the pro-inflammatory, neutrophil recruiting role of TNFα in an acute LPS-induced lung injury mouse model, and its ability to induce endothelial injury in a HUVEC in vitro model.

We studied TNFα signaling in whole blood RNA datasets from human patients with illness ranging from mild respiratory infection to critical illnesses [14, 15]. We observed immunological heterogeneity that was resolved by identifying molecular patient phenotypes with differential enrichment of TNFα pathway activation. As TNFα-inhibitors are major therapeutics in autoimmune disease we assessed whether these phenotypes were shared between critical illness and autoimmune disease.

We leveraged single-cell RNA-sequencing studies of critically ill patients, to identify differences in the cellular composition of individual phenotypes and assess the role of different cell subsets in driving TNFα pathway activation. We propose a novel cell type-based patient classification and associate it with disease severity.

## Methods

### Pre-clinical experiments

All experimental details are provided in the Supplementary materials [16].

### High dimensional-omics datasets

All data was obtained from the public domain. Twelve datasets of whole blood expression profiles from multiple studies of patients with viral respiratory infections, sepsis and autoimmune diseases plus 3 single-cell RNA sequencing (scRNA) datasets from patients with sepsis and COVID-19 were analyzed (Table S1), additionally a bulk proteomics dataset was analysed. Details of the analyzes are provided in the supplementary information (SI).

**Table 1:** Distribution of patient numbers across phenotypes according to disease status.

|  | Sepsis (GSE134347) |  |  | Influenza (GSE101702) |  |  | COVID & Sepsis (COMBAT Consortium) |  |  |  |  |  |
| --- | --- | --- | --- | --- | --- | --- | --- | --- | --- | --- | --- | --- |
|  | Healthy | Non-infectious | Sepsis | Flu mod | Flu severe | Healthy ctrl | HV (Healthy) | COVID Mild | COVID HCW (Health care workers) Mild | COVID Svre | COVID Critical | Sepsis |
| M1 | 0 | 13 | 48 | 14 | 11 | 1 | 2 | 1 | 0 | 1 | 4 | 3 |
| M2 | 0 | 20 | 65 | 6 | 24 | 0 | 0 | 0 | 0 | 0 | 1 | 26 |
| M3 | 0 | 2 | 7 | 3 | 2 | 1 | 0 | 0 | 0 | 2 | 1 | 6 |
| M4 | 0 | 8 | 29 | 29 | 5 | 0 | 0 | 7 | 2 | 18 | 11 | 4 |
| Unc | 83 | 16 | 7 | 11 | 2 | 50 | 8 | 10 | 11 | 20 | 2 | 3 |

### Machine learning to identify phenotypes

A random forest classifier [17] was developed to assign patient samples into phenotypes. Samples were trained against pre-assigned labels from Scicluna *et al*. (Mars1-4) [18], supplemented by a label (“Unc”) to represent healthy controls, using the randomForest and caret R packages [17, 19, 20] (SI). A second classifier was developed using neutrophil cell type GSVA scores and methods were similar to those described above with training on patient clusters identified by their neutrophil subset proportions in the Kwok *et al.* dataset [21].

### Selecting gene features for classification

To generate consistent classification of patients across a spectrum from mild respiratory infection to critical illnesses, we combined gene sets from Scicluna *et al.* [18] with those that distinguish bacterial and viral infections from Sweeney *et al.* [22] and a gene set that identified a pre-vaccination basal inflammatory phenotype [23]. Genes shared with the hallmark TNF pathway gene set were eliminated [24], resulting in 29 genes (Supplementary Table S2).

## Results

We classified patients into molecular phenotypes defined using machine learning across multiple diseases and identified differences in inflammatory signaling specific to these phenotypes. However, we emphasize TNFα pathway signaling in this analysis due to its clinical relevance. We examined the effects of TNFα and TNFα inhibition in an *in vitro* human cellular model and by profiling a mouse lung injury model.

### TNFα drives endothelial damage in human cells and neutrophilia in mouse models of critical infection

We conducted a mouse LPS-challenge study, examining pulmonary responses at various time points: 2-, 9-, 24-, 48-, 72- and 90-hours post-challenge (Fig. 1A). LPS insult induced acute lung inflammation, exemplified by a 2-hour post-challenge peak in bronchoalveolar lavage fluid (BALF) proinflammatory cytokines (including IL-6, KC, and TNFα; Fig. 1C). Subsequently BALF neutrophil levels increased, peaking at 48-hours post challenge and an increase of cellular influx was also observed in the lung via histology (Fig. 1B). We treated mice with anti-TNFα mAb that resulted in a reduction of inflammatory cytokine levels in lung homogenate samples (Fig. 1C). We measured alveolar permeability by examining the concentration of high molecular weight proteins in BALF (Fig. 1D), which increased upon LPS challenge and reversed upon anti-TNFα treatment (Fig. 1D).

**Fig. 1.**
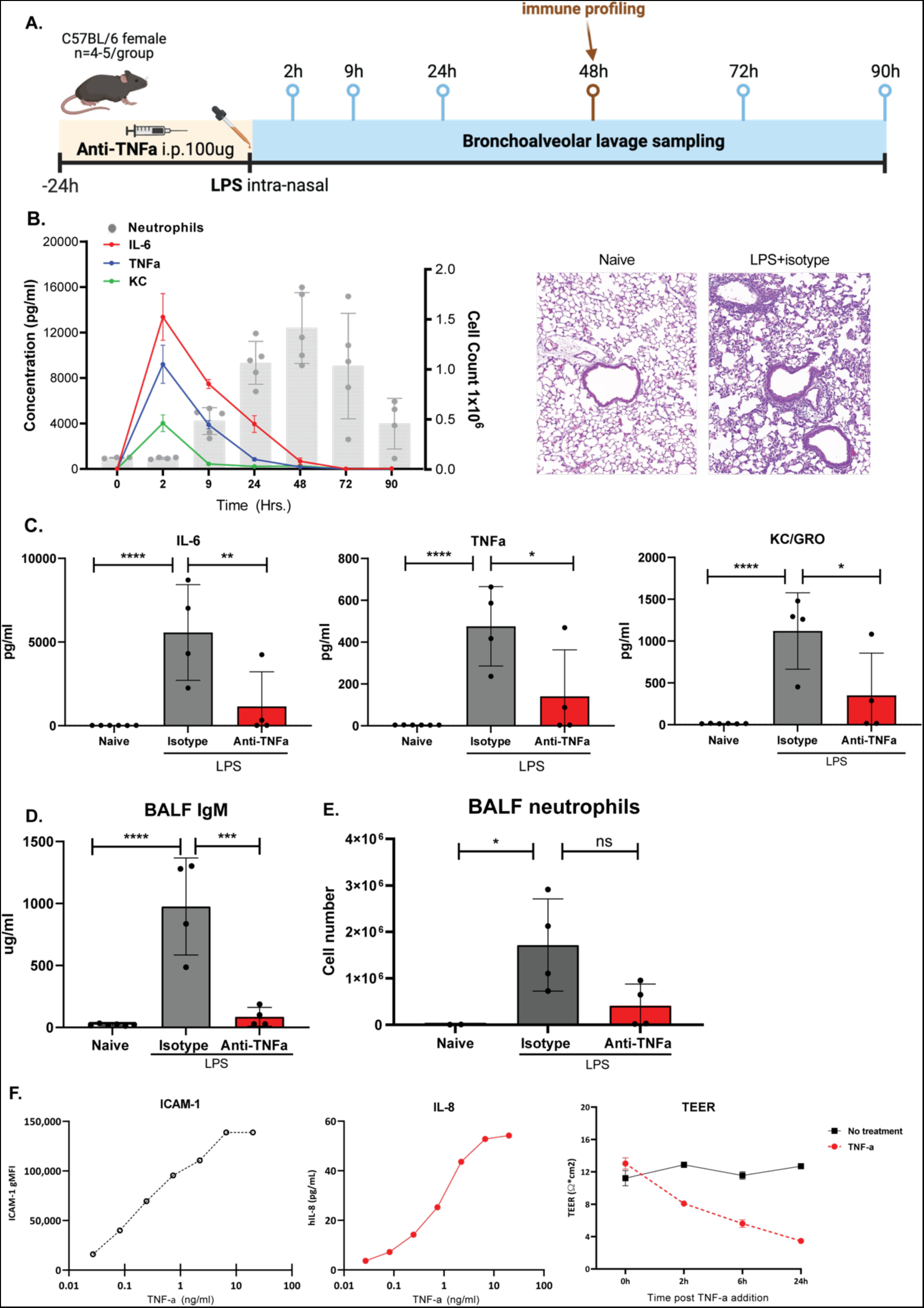
TNF-a is a key driver of pulmonary inflammation in vivo and tissue dysregulation in HUVEC. (**A**) Schematic of LPS time-course mouse experiment. (**B**) Time dependent changes in cell counts in bronchoalveolar lavage fluid (BALF) of mice challenged with LPS. H&E staining of lung session showed tissue damage and immune cell infiltration (**C**) Expression of proinflammatory cytokines in lung homogenate at 48hrs. Mice were treated with 100 ug isotype control or anti-TNFα (**D**) IgM concentration in BALF as a function of LPS challenge and anti-TNFα treatment. **(E)** Neutrophil counts in BALF. Anti-TNFα mAbs reduce neutrophil counts in BALF upon treatment after LPS challenge. **(F)** ICAM-1 and IL-8 concentration in human umbilical vein endothelial cells (HUVEC) across different TNFα concentrations. Trans-Epithelial Electrical Resistance (TEER) after TNFα treatment, shows reduction in barrier resistance at 24hrs. Data are shown as mean ± SEM/SD (n = 1-3).

LPS administration increased BALF neutrophils that reversed upon treatment with anti-TNFα antibodies, which also reduced inflammatory parameters (Fig. 1E). Thus, we show anti-TNFα therapy suppresses inflammation in a mouse acute lung injury model.

To understand how TNFα regulates human endothelial cell function, we utilized human umbilical vein endothelial cells (HUVECs). TNFα induced a dose-dependent increase in HUVEC ICAM-1 and secreted IL-8 as well as a reduced endothelial barrier integrity, discernable by 2 hours after TNFα addition, as measured by reduction in trans-endothelial electrical resistance (TEER) (Fig. 1F).

Given the inflammatory activation in both human in vitro and mouse in vivo models and the effect of TNFα inhibition in the mouse model, we analyzed human transcriptomic datasets to understand the role of inflammation and TNFα signaling across a spectrum of illness.

### TNFα pathway signaling shows a characteristic inflammatory activation and decline in normal infections

To understand immune dysregulation in critical illness, we analyzed the dynamics of immune activation during a normal infection. We hypothesized that critical illness due to infection relates to a dysregulated progression of a normal immune response to infection. In a whole blood dataset from a prospective study of individuals who were followed during and beyond resolution of influenza or rhinovirus infection [25], we found a characteristic initial peak of inflammatory responses, followed by a decline of this response to a subsequent plateau (Fig. 2A-B, Fig. S1A-B). Of the inflammatory signaling pathways, interferon and TNFα activation followed this characteristic pattern, with significant activation within 48 hours of acute symptomatic infection (day 0) relative to baseline (Fig. 2C, pathway normalized enrichment scores (NES) of 1.34 influenza A and 2.14, rhinovirus, respectively and a false discovery rate (FDR) for both pathways < 2.5×10^−02^). TNFα signaling was significantly upregulated in patients with sepsis and pulmonary sepsis relative to corresponding healthy controls (Fig. 2D). This was accompanied by upregulation of other key inflammatory pathways (e.g. IL-6 JAK/STAT signaling). However, TNFα signaling in individual samples showed considerable heterogeneity among patients (Fig. 2D). Consequently, we used machine learning to identify patient phenotypes with consistently enhanced TNFα signaling in critical illnesses.

**Fig. 2.**
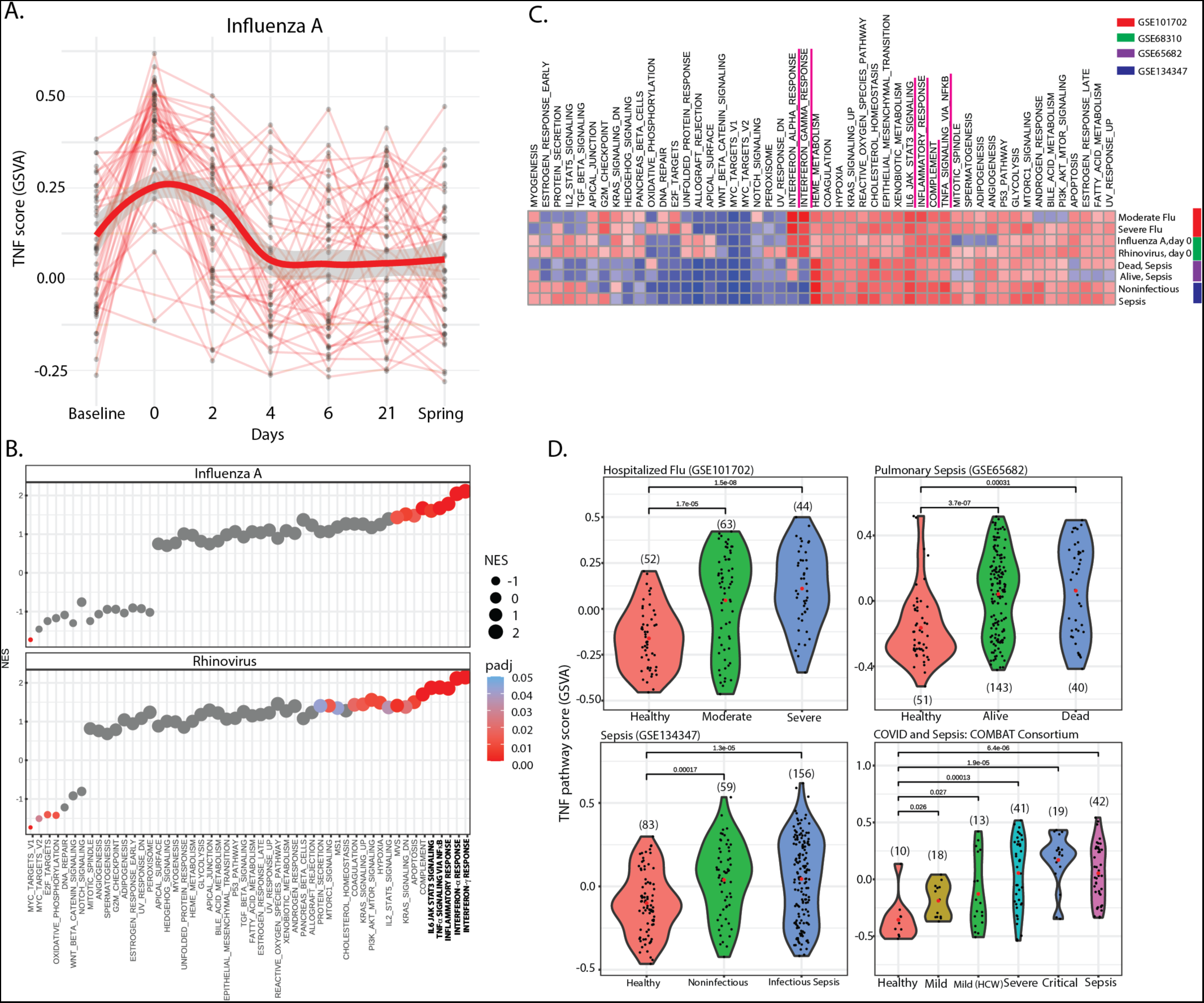
Time-course of immune pathway changes in normal infection versus severe illness. **(A)** Individual trajectories for TNFα pathway activation in whole blood from 64 individuals with Influenza A followed over a flu season (GSE68310), none of whom had severe disease, plotted with a smoothed mean (bright red) and error tolerances (SE) (grey). The trajectories for other viral infections are plotted in Fig S1(A). Data are shown as gene set variation analysis (GSVA) scores for the TNFα pathway module. Infection happens at day 0, baseline is uninfected. **(B)** Dotplots of normalized enrichment scores (NES) and their significance for gene expression differences between Day 0 and Baseline of patients with Influenza A and Rhinovirus infection. The scores and significance are from the gene set enrichment analysis (GSEA) rank test on the (ranked) list of differentially expressed genes (DEGs). There is a strong and significant activation of key inflammatory pathways (highlighted in bold) for both these infections. **(C)** Heatmap of GSEA NES for the hallmark gene set, for DEGs from comparisons of critically ill patients with sepsis (GSE65682: dead sepsis and alive sepsis, GSE134347: sepsis and noninfectious critical illness) or patients hospitalized with Influenza (GSE101702: moderate and severe flu) relative to their healthy controls, plotted alongside scores from comparisons of patients with Influenza A or Rhinovirus at the initial symptomatic day of infection against their healthy baselines (GSE68310) **(D)** Violin plots of TNFα pathway activation (GSVA scores) in individuals from 4 studies (GSE101702, GSE65682 and GSE134347, COMBAT;COVID-19), stratified by disease severity, and healthy controls.

### TNFα signaling is enhanced in a common, inflammatory phenotype that is shared across critical illnesses and is related to neutrophil activation

Previous work on sepsis [18, 26, 27] has shown that clustering patient whole blood expression profiles and assigning phenotypes identifies immunologically homogenous subgroups of patients. We conjectured that phenotyping using the 29 gene signature (Supplementary Table S2) could identify subsets of patients with significantly increased TNFα and inflammatory signaling.

We developed a machine learning classifier to phenotype patients from multiple studies of sepsis, severe influenza and COVID-19 (see methods section)[18, 28–30]. Phenotypes in the training dataset were populated by patients as follows: M1= 22%, M2= 29%, M3= 6%, M4= 16%, and Unc= 27%. TNFα signaling showed consistent activation in patients belonging to the M4 phenotype (Fig 3A). We compared expression profiles of the individual M1-4 phenotypes with control, Unclassified (“Unc”) phenotype. Upon examining TNFα pathway enrichment in each phenotype compared with the reference “Unc” phenotype, or disease state compared to healthy controls, patients in the M4 phenotype consistently showed the greatest, and most significant TNFα pathway activation (Fig 3B) whereas the other 3 phenotypes had more variable associations with TNFα pathway activation across the different disease states. For this reason we focused on the M4 phenotype. The M4 phenotype pathway enrichment profile correlated significantly with pathway level changes seen at the infection visit (day 0) of influenza (severe flu, R = 0.644,p = 3e-07, pulmonary sepsis, R = 0.56, p = 2e-05, sepsis, R = 0.5, p = 1e-04) and rhinovirus patients (severe flu, R = 0.62, p = 8e-07, pulmonary sepsis, R = 0.6, p = 2.5e-08, sepsis, R= 0.61, p= 1.5e-06; Fig. 3C, Fig. S2A, S2B).

**Fig. 3.**
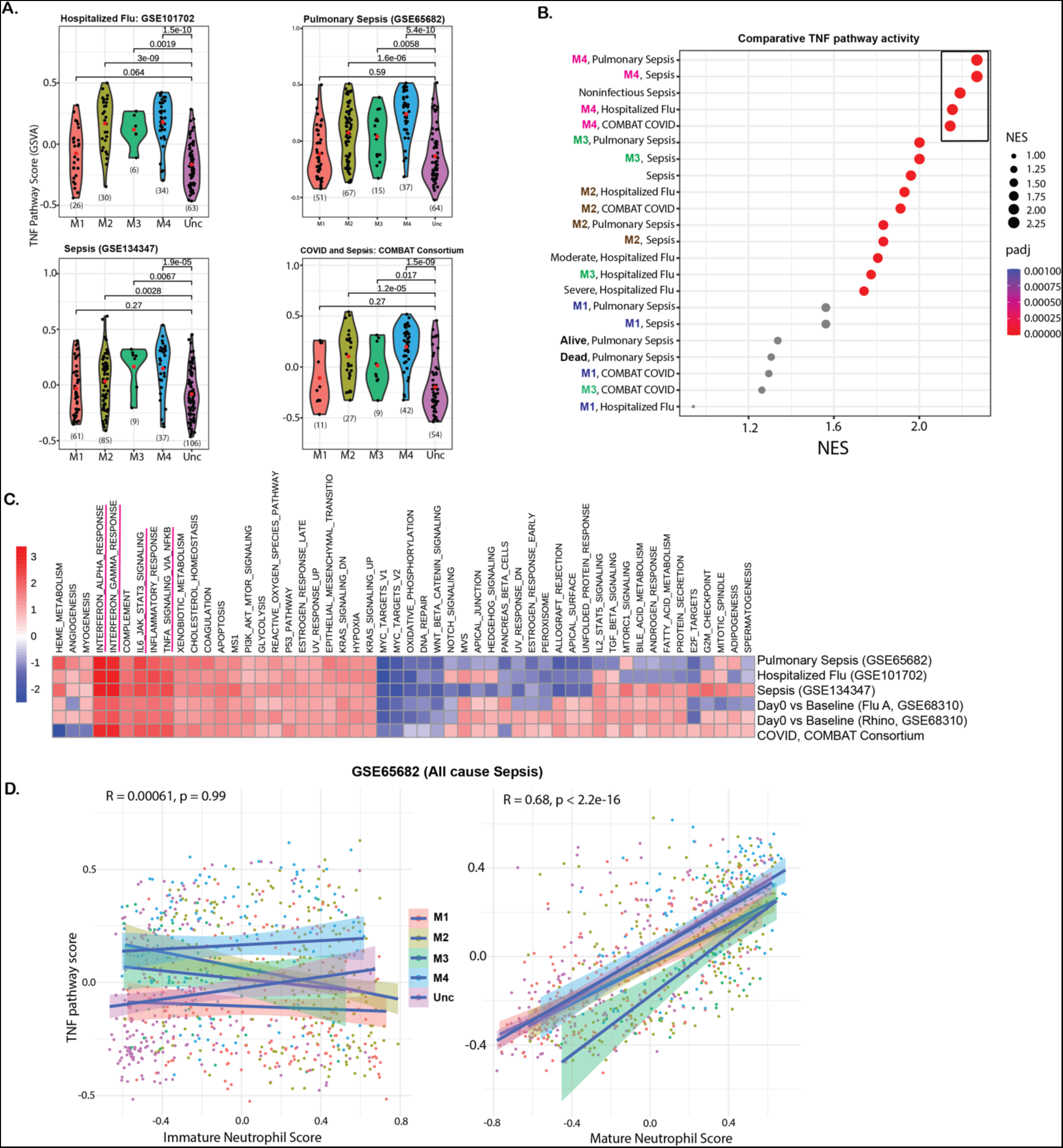
Identifying common phenotypes across infection. **(A)** TNFα pathway profiles of patients distributed into the 5 phenotypes (M1-4 and Unc), show that patients in the M4 phenotype have consistently higher levels of TNFα pathway activation in comparison to the Unclassified (Unc) category (GSVA scores). **(B)** Dotplot of NES for the TNFα pathway across different comparisons. The phenotypes were compared against the reference (“Unc”) phenotype to perform geneset enrichment, while disease states were compared against healthy samples. The most significant and strongly enriched groups are from M4 (inset box). **(C)** Heatmap of TNFα pathway NES estimated from genes differentially expressed between the M4 and Unc categories across multiple studies (GSE68310, GSE65682, GSE134347, GSE101702, COMBAT). The pathway profile of M4 patients is remarkably consistent between studies and different forms of infectious illness. **(D)** Scatter plot of TNFα pathway activation with immature and mature neutrophil signature score for patients with Sepsis (GSE65682).

The M4 phenotype represented a conserved pattern of immune activation and associated with upregulated interferon signaling as well as IL6-JAK/STAT pathway activation (Fig. 3C, S2A).

We generated a gene signature (using Wilcoxon differential expression) that distinguishes the differently annotated neutrophil subsets identified in published single-cell RNA-sequencing (scRNA) datasets [31] (Supplementary Data 1), and observed that enrichment of the mature, but not immature, neutrophil signature strongly correlated with TNFα pathway enrichment in whole blood from patients with all cause sepsis (Fig. 3D). Our analysis identified a high inflammatory subset of patients (M4), with increased Interferon (3.2 > NES > 3.01, FDR < 2e-40 in severe illness) and TNFα signaling (2.24 > NES > 1.96, FDR < 4.0e-07). TNFα pathway activation correlates with enrichment of mature neutrophil genes in whole blood from these patients.

anti-TNFα inhibitors are a mainstay of treatments for auto-immune disease, hence we examined whether our phenotypes extend to auto-immune disease and similarly correlate with TNFα signaling. Moreover, the link between systemic immune changes and localized tissue injury can be investigated in autoimmune disease patients, while impractical in critically ill patients. Thus, a link with autoimmune disease could enable inference of tissue injury mechanisms in critical illness.

### A subset of patients with autoimmune disease belong to the inflammatory sepsis phenotype

We investigated the presence of a shared immune phenotype between chronic autoimmune diseases and critical illnesses, and furthermore whether TNFα activation was correlated with neutrophil content. The interferon-associated M4 phenotype was seen across autoimmune diseases with an inflammatory pathway profile like M4 patients with sepsis and respiratory illness (Fig. 4A, supplementary figure S3) [32–35]. In a dataset from lupus patients [36], TNFα pathway scores correlated strongly and highly significantly with neutrophil percentages (Fig. 4B). Among the phenotypes, patients with the M4 phenotype had a higher neutrophil percentage (Fig. 4B). In another Lupus dataset [35], for which clinical ISM (Interferon signature metric) scores [35, 37] were available, the M4 phenotype consisted entirely of patients with high ISM scores (i.e., high interferon activation, Fig. 4C). We found a strikingly high correlation between TNFα pathway scores with mature neutrophil marker scores across datasets of patients from ulcerative colitis, rheumatoid arthritis and lupus studies (Fig. 4D).

**Fig. 4.**
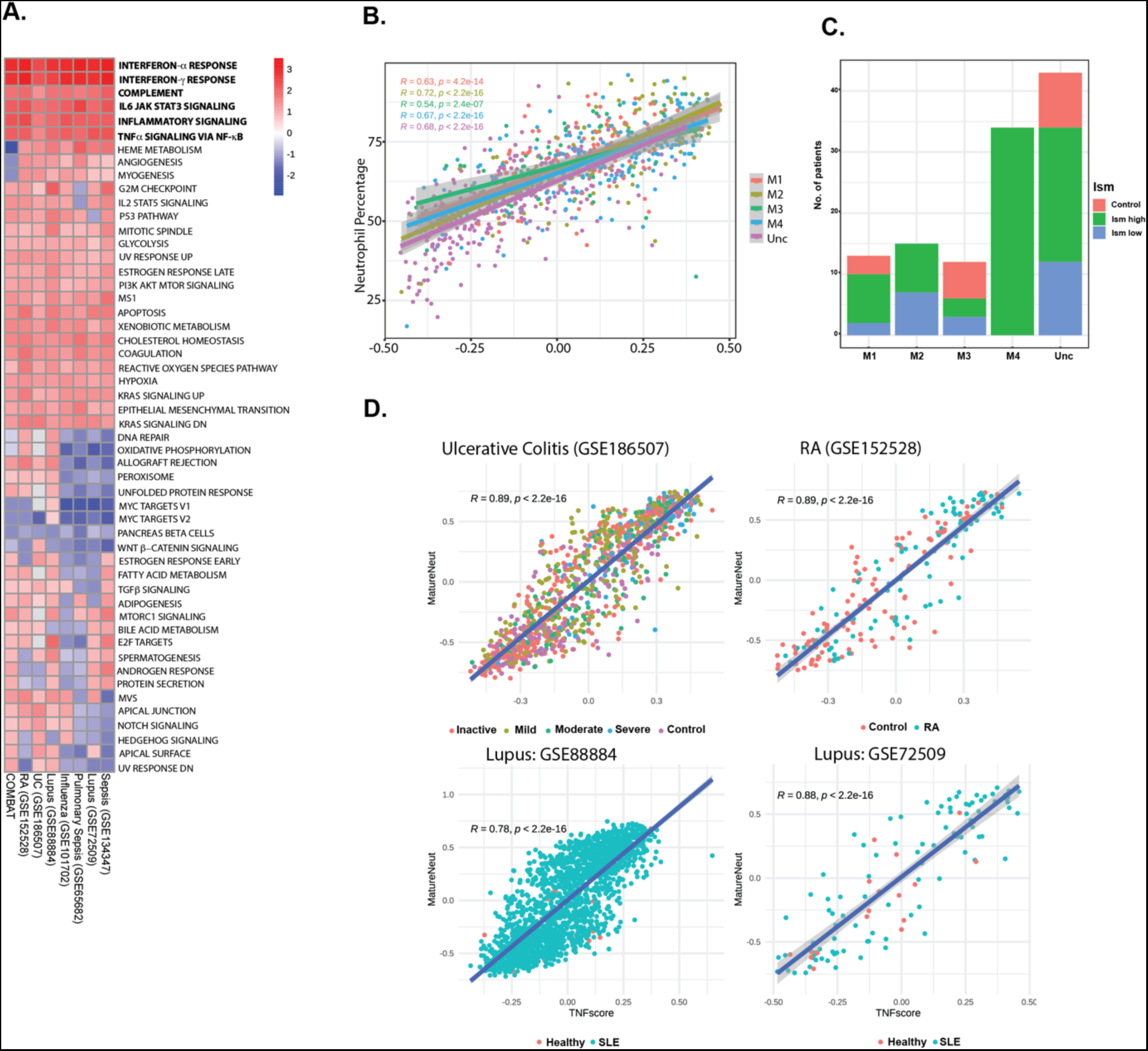
TNF signaling and phenotype stratification in autoimmune disease. **(A)** Heatmap of GSEA NES for the Hallmark pathway gene set, derived from a differential gene expression analysis of patients with autoimmune diseases categorized into M4 compared against a corresponding unclassified (Unc) phenotype (GSE88884, GSE72509, GSE186507, GSE216678, GSE101702, GSE65682, GSE134347, COMBAT). The heatmap makes clear the common patterns of inflammatory activation across diseases in this phenotype. **(B)** Measured neutrophil percentages are plotted against corresponding TNFα pathway activation scores in Lupus patients (GSE65391). We note a strong correlation between the percentage of neutrophils in the blood and TNFα activation, with patients in M4 having an overall higher neutrophil percentage. **(C)** Distribution of ISM status in lupus patients and healthy controls across the M phenotypes (GSE72509). The M4 phenotype is entirely composed of ISM high Lupus patients. (**D**) Scatterplots of TNFα enrichment scores from GSVA against corresponding scores for mature neutrophil marker gene sets for datasets from multiple autoimmune conditions.

Taken together, our analysis established a high inflammatory phenotype that strongly correlates with increased TNFα pathway activation in patients hospitalized with infection. Furthermore, TNFα activation correlates most strongly with neutrophils in whole blood, and anti-TNFα therapy resolved neutrophilia and inflammation in a mouse model of acute lung injury. Thus, we leveraged publicly available scRNA data from sepsis patients to better understand the cellular origins of TNFα pathway and phenotype heterogeneity in sepsis.

### Single-cell RNA-sequencing analysis shows that sepsis phenotypes have an altered neutrophil composition

We analyzed two COVID-19 scRNA studies (COMBAT UK consortium PBMC and whole blood [38]) and a German COVID-19 consortium PBMC and neutrophils [31]) and a whole blood scRNA study of sepsis patients and healthy controls [21]. Phenotypes in the UK COVID study were estimated using whole blood RNA data from patients with scRNA profiles, while the other two studies were predicted on sample level pseudo-bulking of scRNA counts (Methods). After prediction in these studies, the M4 phenotype consisted of patients with severe and critical illness (Table S4), while the M2 phenotype was enriched for sepsis patients. We found no significant difference in abundances of cells from the PBMC compartment from the COMBAT dataset between TNFα pathway high and TNFα pathway low patients, or between the M4 and Unc phenotypes. However, TNFα signaling pathway enrichment was observed for many innate immune cell types in the M4 phenotype (Supplementary Figure S4A). TNFα signaling correlated with mature neutrophil abundances, with the M2 phenotype being enriched for immature neutrophil populations unique to sepsis patients (Fig. 3D, 4D, 5A).

**Fig. 5.**
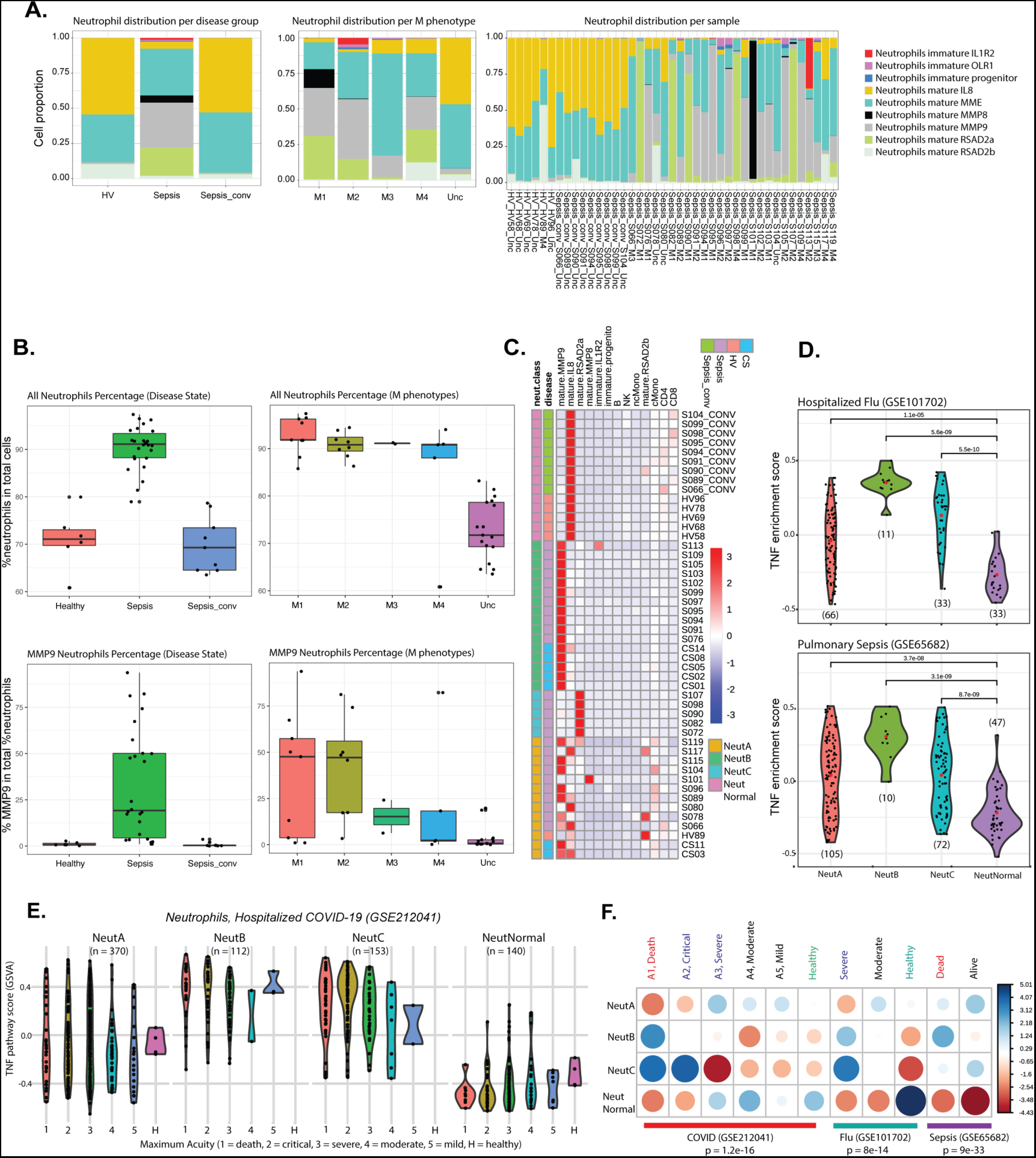
Cell subpopulations associated with phenotypes and increased TNF signaling. **(A)** Neutrophil subset proportions as a function of disease, phenotype and individual patient samples (Kwok Sepsis dataset). **(B)** Distributions of neutrophil total and MMP9 frequencies across phenotypes (Kwok Sepsis dataset). **(C)** Heatmap of abundances of neutrophil subsets identified from scRNA analysis. The heatmap shows the existence of at least 4 patient subtypes (Kwok Sepsis dataset). **(D)** Distribution of GSVA scores for the TNF signaling pathway from phenotyping on neutrophil subset enrichment scores. The phenotype model was trained on the clustering shown in (**C**). **(E)** GSVA enrichment scores for the TNFα signaling pathway for each neutrophil phenotype, grouped by disease (COVID-19) severity. Severity decreases from class 1 to 5, with 1 representing 28-day mortality. The data here is from bulk RNA profiling of sorted neutrophils. **(F)** Pearson residuals of patient populations distributed across phenotypes and disease status. The more positive (blue) residuals correspond to a greater than expected number of patients in a given phenotype and disease category. The size of the circles correspond to the relative proportion of patients.

We analyzed neutrophil profiles from the whole blood scRNA dataset of sepsis patients (Kwok *et al.* [21]). By grouping neutrophils with an iterative clustering approach, we were able to identify 3 immature subtypes of neutrophils: immature progenitor (*DEFA3, AZU1, MPO, ELANE*), immature OLR1 (*LTF, OLR1, CEACAM8*) and immature IL1R2 (*LTF, SELL, IL1R2, CD44*) (Supplementary Figure S4B). On the mature compartment, 6 subtypes could be identified: mature IL8 (*CXCR2, CXCL8, CHI3L1*), mature MMP8 (*CXCR2, MMP8, S100A12*), mature MMP9 (CXCR2, CD177, MMP9, ARG1), mature MME (*CXCR2, MME*), mature RSAD2a (*CXCR2, IFI6, RSAD2, SLAMF7*), mature RSAD2b (*RSAD2, MX1, ISG15*) (Supplementary Figure S4C). Interestingly, sepsis donors contained the different immature subtypes, together with mature MMP8 and MMP9 (Fig. 5A). After sepsis convalescence, donors showed a neutrophil compartment mainly constituted by mature IL8, MME and RSAD2b, with comparable cell composition of healthy donors (Fig. 5B). Compositional comparison across M classification phenotypes, showed that the mature MMP8 population is characteristic of M1, whereas M2 is characterized by immature neutrophils (Fig. 5B).

This dataset showed a higher percentage of neutrophils in whole blood samples of sepsis, compared with healthy donors and sepsis patients after convalescence (Fig. 5B). The unclassified (“Unc”) samples also showed lower percentage of neutrophils in comparison to patients in M1-M4. The mature MMP9 neutrophil percentage in total neutrophils showed a good separation between M1 and M2 (Fig. 5B) and sepsis. Neutrophil diversity estimates from a Shannon index score calculation showed a lower average diversity in neutrophils in healthy individuals, versus sepsis and sepsis after convalescence (Fig. S4F). Moreover, a lower average diversity could be observed for M1 (Fig. S4F). Additionally, richness of neutrophil estimates was higher in sepsis when compared with healthy and higher trend in M1 and M2 (Supplementary Figure S4F). These observations suggest that our phenotypes can be partially explained by their neutrophil content. This motivated us to stratify patients based on their neutrophil content and study the properties of the resulting phenotypes.

### Sepsis patients segregate into subgroups with different neutrophil composition

The distribution of neutrophil subpopulations segregated samples in this dataset [18] into 4 clusters (NeutA, NeutB, NeutC and NeutNormal, Fig. 5C). Specifically, NeutNormal patients have high mature IL8 neutrophil composition, while NeutB have high mature MMP9 neutrophils, NeutC is composed of high RSAD2 neutrophils and NeutNormal contains a mixed population (Fig 5C). We generated marker gene signatures (Supplementary Data 1) for each neutrophil subpopulation and estimated their gene set enrichment (GSVA) scores in patient pseudobulk profiles and constructed a random forest model to stratify patients into the 4 clusters identified from clustering their population percentages (Fig. 5C). Application of the cell type dependent classifier to the other datasets of critical illness (Table S5) showed that TNFα pathway signaling is significantly higher in the NeutB, and to a lesser extent in the NeutC phenotypes compared to NeutNormal (Fig. 5D). On phenotyping hospitalized COVID-19 patients based on bulk RNA profiles of neutrophils from whole blood (GSE212041), we find upregulation of TNFα pathway signaling in the NeutB and NeutC phenotypes, and furthermore TNFα pathway signaling consistently, but not statistically significant, increases with severity of illness within these phenotypes (Fig 5E).

We assessed the correlation between membership in different phenotypes with disease severity or mortality, in 3 datasets where this information is available (GSE212041, GSE65682, GSE101702). We find that membership in either of NeutB or NeutC phenotypes correlates with disease severity, with NeutB and NeutC patients having significantly more severe disease (Fig. 5F). Patients in NeutB have a mortality of 50%, in comparison with mortality rates of about 20% in NeutA and NeutC in the sepsis (GSE65682) cohort, although this may not be significant due to the small number of patients in NeutB (10 patients).

## Discussion

We tested anti-TNFα mAb as a therapy in a mouse model of ARDS and examined its effect on lung injury induced by LPS stimulation. We found that anti-TNFα therapy reduced LPS-induced lung inflammation including neutrophilia and enhanced alveolar permeability.

We used machine learning to stratify patients with sepsis and severe infection into homogenous immune phenotypes. We identified patients with a common high inflammatory phenotype, wherein TNFα signaling was significantly more activated. We found patients with this phenotype upon expanding our phenotype classification to studies of multiple autoimmune diseases. TNFα signaling was correlated with mature neutrophil levels in severe and critical illness, which also extended to autoimmune diseases. This is consistent with previous reports that anti-TNFα therapy reduced neutrophil content in patients with autoimmune diseases [39–43]. Both our mouse studies and the computational analysis of human data show activation of additional inflammatory pathways, notably the IL-6 and IFN signaling pathways, which suggest avenues for additional targets in sepsis and ARDS.

Our work shows the prevalence of neutrophil subsets in sepsis patients that correlate with disease severity in respiratory illness implying that granulopoiesis is altered, in agreement with previous studies [21, 44]. Interestingly our work suggests that much of the heterogeneity in patients hospitalized with respiratory infection is due to differences in specific neutrophil subsets, implying that granulopoiesis is heterogeneously altered. Correspondingly, we find that none of the other major cell subsets show significant change in abundances for the phenotype classifications that we analyzed. Indeed, a classifier based on neutrophil subsets was able to clearly stratify patients and the resulting phenotypes correlated with disease severity. This does not preclude changes in the functional state of different cell types, and indeed single cell studies of the PBMC compartment have identified monocyte derived MDSCs in severe COVID and sepsis [45, 46]. Together with our work, this suggests the critically ill individuals show a broad shift in immune cell development.

Clinical studies have shown remarkable efficacy for anti-TNFα therapies in multiple auto-immune inflammatory diseases. However, anti-TNF therapies have shown modest success in reducing mortality in critical illness. The ACTIV-1 trial of infliximab in patients hospitalized with COVID-19 (n=518) showed a substantial improvement in mortality in the treated group (10%) compared to placebo (14.5%), while a meta-analysis of 17 clinical trials in sepsis showed a significant reduction in all-cause mortality in patients treated with anti-TNFα mAbs (OR = 0.91, p = 0.04) and a trend towards better survival for patients with high IL6 levels (>1000 pg/ml, OR = 0.85) upon anti-TNFα treatment [10]. Notably, the RAMSES trial of sepsis patients stratified by IL-6 levels, treated with afelimomab or placebo was terminated early for lack of efficacy [47].

These studies suggest that anti-TNFα treatment could have efficacy in critically ill patients who have phenotypes with hyperinflammatory activation. An advantage of identifying a universal, hyperinflammatory phenotype that is prevalent in both acute and chronically inflammatory states is that systemic changes in immune activation can be correlated with localized mechanisms of tissue injury in chronic autoimmune disease, for which tissue is available. This correlation can be used to infer similar mechanisms in critically ill individuals allowing for novel therapies and precision medicine approaches to treat critical illness.

### Limitations

Although our work has identified phenotypes of patients who have significantly enriched TNFα signaling, we cannot directly assess the impact of anti-TNFα therapy on these patients due to unavailability of study data in this regard. Our analysis relies on human whole blood transcriptomics data, however the correlation between systemic changes in the immune response during critical illness with pathological processes driving local tissue injury is not well understood. Thus, although we can classify patients into phenotypes, these may remain heterogeneous in the pathological mechanisms of tissue injury and could impact stratification strategies that build on systemic immune response-based phenotyping.

Our work infers neutrophil subset proportions by proxy with enrichment scores of corresponding marker signatures, since information about abundances of neutrophils and their subsets is usually unavailable because most datasets are of PBMCs. This is a weakness in our work since enrichment scores and neutrophil cell subset abundances may differ.

### Future Directions

While our work identifies biologically homogenous phenotypes (“predictive enrichment”), patient phenotypes that are homogenous in their clinical presentation (“prognostic enrichment”) have been identified through the application of machine learning to clinical data in ARDS and Sepsis [4, 5]. An interesting future direction of research is to investigate the relationship between these two approaches that would rely on a joint analysis of clinical and molecular or biological properties. This could inform precision medicine treatments of critical illness.

## Supplementary Information

### Supplementary Methods

#### Ethical Statement and Animal Experimentation

All experimental animals used in this study were under a protocol approved by the Institutional Animal Care and Use Committee of Janssen Pharmaceuticals, Johnson & Johnson. The mice were housed in disposable cages (Innovive) and received food and water ad libitum. Before experimentation, the mice were allowed to adapt to the experimental environment for a minimum of one week.

The criteria of the American Chemical Society ethical guidelines for the publication of research were met. Every effort was made to minimize animal discomfort and to limit the number of animals used. Mice were kept in a specific pathogen-free facility under appropriate biosafety level following institutional guidelines.

#### LPS induced acute lung injury model

6-8 weeks female C57BL6 mice (JAX #0006664) were used. Mice were anesthetized by isoflurane. E. coli-derived LPS (O111.B4, Sigma-Aldrich L4191-1MG) was administered intranasally (10 μg/mouse in 35 μl). Mice were sacrificed at 2, 9, 24, 48, 72, 90 hrs post LPS insult. Lung and BALF were collected following standard procedure (Previously described by Rao and colleagues: PubMed 20110560).

#### LPS induced acute lung injury model treated anti-TNF-α

6-8 weeks female C57BL6 mice (JAX #0006664) were used. Mice were anesthetized by isoflurane. E. coli-derived LPS (O111.B4, Sigma-Aldrich L4191-1MG) was administered intranasally (10 μg/mouse in 35 μl). Mice were sacrificed at 48 hrs post LPS insult. Lung and BALF were collected following standard procedure. Two doses of anti-TNF-α of 100 μg per animal (BioXCell #BE0058) and isotype control (BioXCell #BE0088) were given i.p. 24 hours and 1 hour before LPS insult.

#### Cytokine quantification

BALF and lung cyto-, chemo-kines were assessed using the Proinflammatory Panel 1 and Cytokine Panel 1 kits for mice (Meso Scale Diagnostics) with results collected on a MESO QuickPlex (Meso Scale Diagnostics) and analyzed with Discovery Workbench 4.0 (Meso Scale Diagnostics), per manufacturer’s directions. ELISA BALF IgM level was measured by the Mouse/Rat IgM ELISA kit (abcam, ab215085) following the manufacturer’s instruction. IL-8 levels were detected in the collected HUVEC culture supernatant at 4h and 24h using human IL-/CXCL8 DuoSet ELISA (R&D Systems DY208) following manufacturer’s instructions.

#### Flow cytometry

For extracellular antigens, freshly isolated cells were stained with antibodies for 20 minutes at 4 °C in FACS buffer (PBS + 0.5% BSA), then washed and acquired by flow cytometry. Cells were resuspended in FACS buffer before data acquisition. Cells were collected on BD LSRFortessa or FACSymphony. Data was acquired using FACSDiva (BD) and analyzed with FlowJo (BD Biosciences) software. Flow cytometry antibody panels and reagents used can be found in the additional table and gating strategy (Fig. S6).

#### Antibodies for flow cytometry

| Antigen | Fluorochrome | Clone | Vendor | Cat# |
| --- | --- | --- | --- | --- |
| Human CD54/ICAM1 | PE | HDC54 | BioLegend | 322708 |
| Human CD31 | APC | WM59 | BioLegend | 303116 |
| Mouse CD45 | AF700 | 30-F11 | BioLegend | 116126 |
| Mouse CD11b | BUV395 | M1/70 | BD Biosciences | 563553 |
| Mouse Ly6G | PE-Dazzle594 | 1A8 | BioLegend | 127647 |
| Live/dead | Yellow |  | eBioscience | L34959 |
| 7AAD (Cell viability) |  |  | eBioscience | 00-6993-50 |

#### TNF-α stimulation of Human Umbilical Vein Endothelial Cells (HUVECs)

HUVECs (Lonza C2517A, donor 22TL018260) were seeded in a 24-well TC-treated plate utilizing EGM2 (Lonza) media. 48h post seeding, recombinant human TNF-α (R&D Systems 210-TA-020) was added at different concentrations, 20 ng/mL diluted down 3-fold to 27 pg/mL with PBS. Cells were incubated for up to 24h. Supernatant was collected at 4h and 24h to check for IL-8. To measure levels of ICAM1, cells were detached (Lonza CC-5034) and stained immediately with antibodies for 20 minutes at 4°C in PBS, washed, resuspended in PBS before data acquisition on a Bectin Dickenson (BD) Fortessa flow cytometer.

#### Choice of human transcriptomic datasets

We chose datasets for analysis according to the following criteria: a) they should be whole blood mRNA profiles. We made an exception for a dataset of neutrophil profiles for reasons detailed in the manuscript. b) they should correspond to moderate (n > 100) to large sample sizes. We relaxed this criterion somewhat for scRNA datasets to be considered, and for time course studies. c) they should together represent a broad spectrum of illness. We list the datasets considered below.

The total number of samples covered by these data is n = 7,886 corresponding to approximately 6,000 individuals.

**Table S1:**
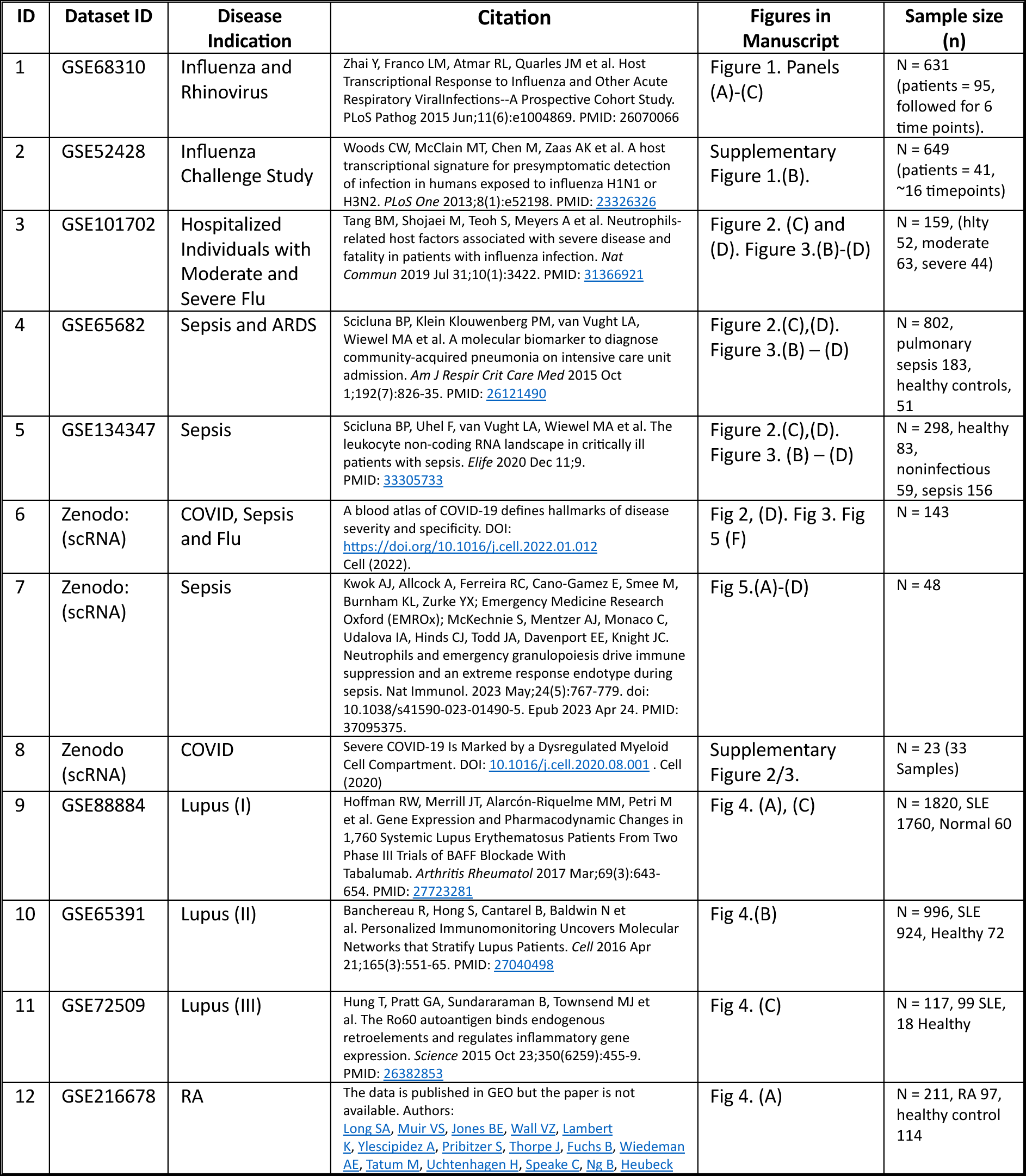
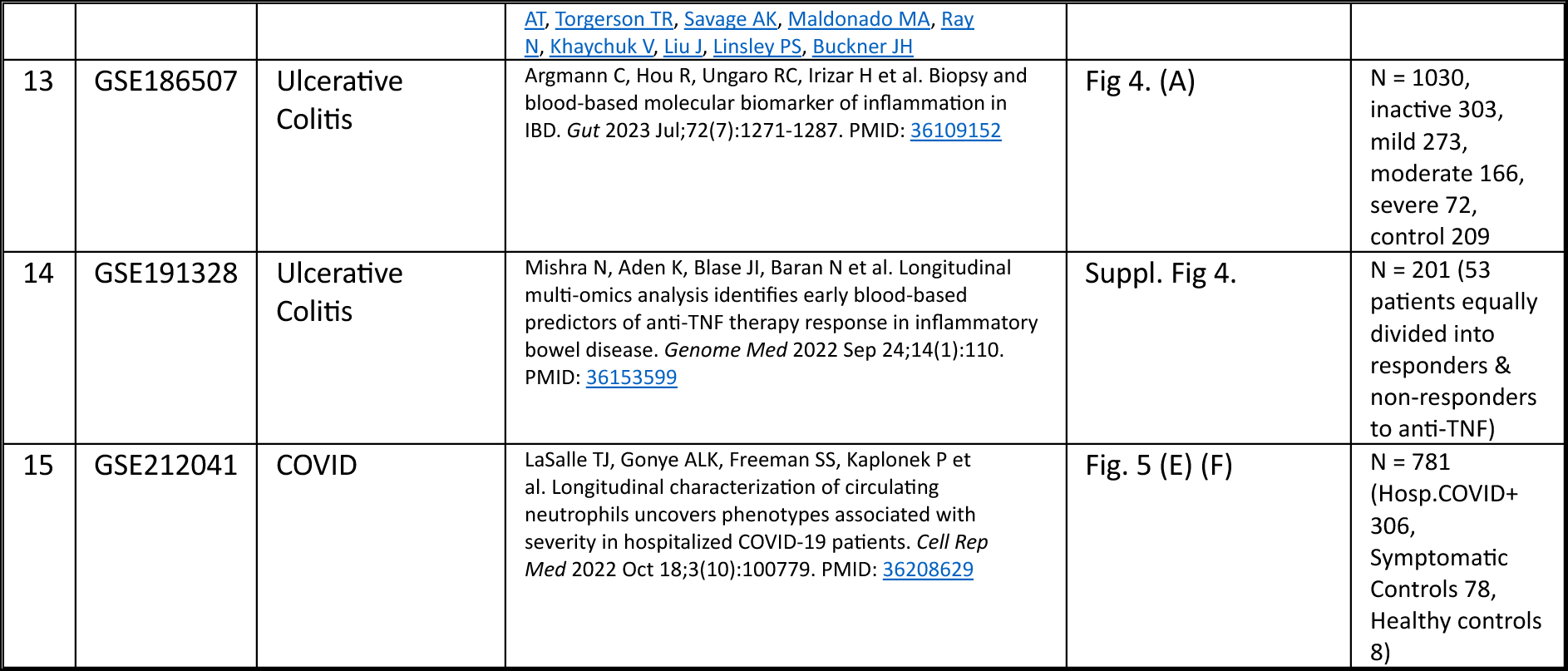
Datasets Analyzed in this Study.

#### Bulk RNA-seq and data analysis

For the whole blood expression microarray datasets, processed log expression values at the probe level, along with their annotations were downloaded from NCBI GEO, and downstream analysis performed, while for RNA sequencing data, raw counts were downloaded along with relevant metadata. For the RNA counts datasets, variance stabilizing transformation was used to convert them to normalized log expression values for use in clustering, phenotype classification and estimating individual sample level pathway enrichment. Processed microarray expression values were used as is. Normalization and differential expression analysis for the RNA counts data was performed using edgeR [15–17], and limma [18] was used for this task on the microarray datasets. Pathway analysis was performed on the Hallmark pathway gene set collection [19] using an implementation of the Gene Set Enrichment Analysis (GSEA) method [20, 21] in the fgsea R package [22], and sample specific enrichment through the Gene Set Variation Analysis (GSVA) method [23].

#### Single-cell RNA-seq and data analysis

The single-cell RNA-seq dataset from Kwok *et al.* [24] was used for a more in-depth analysis. Pre-processed data along with their annotations were downloaded from Zenodo (DOI: 10.5281/zenodo.7723202). All downstream analysis was done in R using the Seurat v4.1.1 package. Raw count data was log-normalized with a scaling factor of 10,000. For the first level clustering, the top 1,500 most variable genes were selected (‘vst’ method implemented in FindVariableFeatures()) and scaled the ScaleData() function. Afterwards, principal component analysis (PCA) was run, the number of significant principal components (PCs) to be used for downstream cell clustering was determined using an ElbowPlot and heatmap inspection. A nearest neighbour graph and Uniform Manifold Approximation and Projection (UMAP) plot were generated using the significant PCs, followed by louvain clustering, and the best resolution for clustering was determined using an average silhouette score approach, testing 10 resolutions between 0.1 and 1 as previously described in Ziegler et *al.* [25]. Marker genes for each cluster were calculated using the FindAllMarkers() function (method= ‘wilcox’) and each cluster was iteratively subclustered further using the same approach. Sub-clustering was stopped when the resulting clusters were not meaningfully different.

A subpopulation pseudo-bulk aggregation was performed using muscat v1.4.0 [26]. Each sample neutrophil raw data counts were aggregated to pseudo-bulk data using the aggregateData function with the fun=“sum” option. Downstream Hallmark 50 pathway analysis was performed with an implementation of the Gene Set Enrichment Analysis (GSEA) method in the fgsea R package, and sample specific enrichment through the Gene Set Variation Analysis (GSVA) method. Richness and diversity metrics of neutrophils per sample were calculated in R using the vegan package.

#### Developing subphenotype classifiers

##### a. Gene signature selection

We used a gene set for the M classifier as listed below.

**Table S2:** Genes used to develop the M classifier.

|  |  |  |  |  |  |
| --- | --- | --- | --- | --- | --- |
| BPGM | PDCD10 | IFI27 | SERPING1 | RSAD2 | OAS1 |
| TAP2 | IFIT5 | JUP | ANXA3 | DHX58 | OAS3 |
| GADD45A | GLTSCR2 | LAX1 | ELANE | FCER1A | PML |
| PCGF5 | HK3 | IRF7 | C1QB | HERC5 | CXCL16 |
| AHNAK | CTSB | RARA | DDX50 | IFIT1 |  |

##### b. Developing a random forest classifier

To develop our phenotyping classifiers, we used the random forest method as implemented in the randomForest R package. We used the following parameters: mtry = 5 (for the M subphenotype classifier) and mtry = 3 for the neutrophil classifier. We used ntree = 1500 for both. The relevant code and R scripts are deposited along with this document.

We tested different approaches to perform this classification: first we used the log normalized RNA expression values as is, and we found that this performs poorly due to the highly variable range of expression values demonstrated by different features in the classifier. We thus reasoned that using a z-score transformation of the values for each feature that preserves their relative ordering while scaling their values to have a comparable range across features would be a better choice. Our final classification uses this z-score transformation. The specific transformation used is defined below:

Let 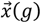 define the expression profile of a gene across the samples in a dataset (each element of the vector is the log expression value of the gene for a given sample). Then, we use the following transformation:

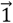

Here |*S*| is the number of samples in the dataset *S*, 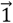 is the vector of all 1’s and 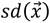 is the standard deviation of the gene expression values across the samples *S*.

For datasets that had RNA seq count values, we tested using the log normalized counts (log cpm) values, as well as the values derived from a variance stabilizing transformation (vst). We found that there are differences in the classification using the two types of normalized counts, but eventually decided on using the vst normalized counts, since we reasoned that it is appropriate to account for dispersion effects both for our single sample pathway analysis using GSVA, and consistently, for the purposes of phenotype classification. We note that similar differences showed up when we tested the SRS phenotyping classifier for such data.

We trained our gene-based classifier against the “Mars” labels defined by the MARS consortium. A table of our predicted class and the MARS labels for pulmonary sepsis patients from GSE65682 is shown below.

**Table S3:** Patient numbers across M phenotypes vs Mars phenotypes: GSE65682.

|  | Mars1 | Mars2 | Mars3 | Mars4 | NA |
| --- | --- | --- | --- | --- | --- |
| **M1** | 37 | 4 | 3 | 3 | 4 |
| **M2** | 2 | 56 | 7 | 0 | 2 |
| **M3** | 0 | 1 | 14 | 0 | 0 |
| **M4** | 5 | 3 | 9 | 20 | 0 |
| **Unc** | 0 | 0 | 18 | 1 | 45 |

Similarly, we show the relation between disease status and our predicted M phenotypes below across different critical illness datasets that we have analyzed:

**Table S4:**
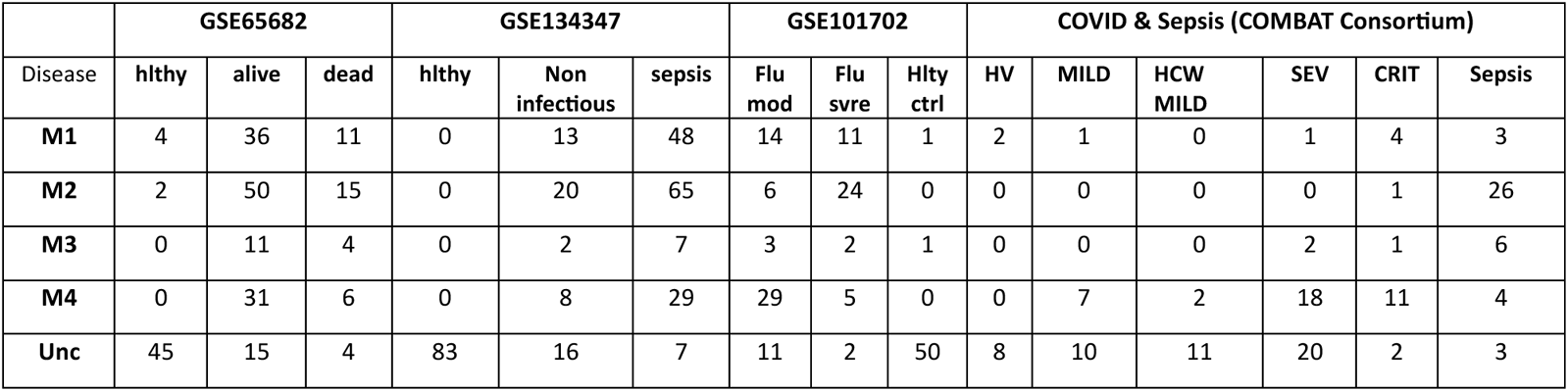
Patient numbers across M phenotypes vs Disease.

We also applied the neutrophil classifier across these and an additional covid dataset. The results are shown in Supplementary Table 5.

**Table S5:** Patient number across Neutrophil phenotypes vs Disease.

| GSE212041 (COVID-19) | GSE101702 (Flu) | GSE65682 (Sepsis) |
| --- | --- | --- |

| condition | 1<br>(death) | 2<br>(Critical) | 3<br>(Severe) | 4 | 5 | H | Flu<br>svre | Flu<br>mod | Hlty<br>ctrl | 0 | 1<br>(death) | H |
| --- | --- | --- | --- | --- | --- | --- | --- | --- | --- | --- | --- | --- |
| NeutA | 31 | 97 | 181 | 37 | 19 | 4 | 17 | 44 | 32 | 79 | 21 | 5 |
| NeutB | 27 | 34 | 45 | 2 | 3 | 0 | 7 | 5 | 0 | 5 | 5 | 0 |
| NeutC | 38 | 75 | 33 | 7 | 2 | 0 | 20 | 13 | 0 | 54 | 14 | 4 |
| Neut<br>Normal | 8 | 31 | 75 | 21 | 7 | 4 | 0 | 1 | 20 | 5 | 0 | 42 |
| **COVID & Sepsis (COMBAT Consortium)** |  |  |  |  |  |  |  |  |  |  |  |  |
|  | Sepsis |  | CRIT |  | SEV |  | HCW<br>MILD |  | MILD |  | HV |  |
| NeutA | 15 |  | 10 |  | 28 |  | 4 |  | 7 |  | 2 |  |
| NeutB | 5 |  | 2 |  | 3 |  | 0 |  | 1 |  | 0 |  |
| NeutC | 22 |  | 7 |  | 4 |  | 0 |  | 0 |  | 0 |  |
| Neut<br>Normal | 0 |  | 0 |  | 6 |  | 9 |  | 10 |  | 8 |  |

##### c. Mature and Immature Neutrophil Gene Signatures

To assess the correlation of TNF signaling with neutrophil levels, and also the gene sets that were used to train the neutrophil based cell classifier, we used neutrophil signatures as described in the Supplementary Data (excel sheet tab “NeutrophilSignatures”).

#### Differential and Pathway Enrichment Analysis

For microarray expression data, we downloaded normalized expression values from ncbi geo using their GEOquery R package. We performed differential expression analysis using the limma-empirical Bayes framework, and pathway enrichment on differentially expressed genes using the fgsea package. Similarly, for RNAseq counts data, we obtained raw counts, filtered out low or no-expressing genes using edgeR’s filterbyExpr framework, typically using a design matrix constructed from the disease phenotypes of the samples. Subsequent to this, we performed library normalization, dispersion estimation and differential gene expression using edgeR’s framework with glmQLFit and glmQLFTest functions.

For single sample pathway enrichment, we used the GSVA package from Bioconductor, which estimates single sample gene set enrichment after a z-score like transformation of the data. We used these pathway enrichment scores to estimate patient TNF, Interferon and other pathway activation status, and also to perform neutrophil content based subphenotyping. We resorted to using neutrophil marker enrichment scores as a proxy for neutrophil cell proportions since we usually did not have data regarding neutrophil and neutrophil subset composition in most studies that we considered.

#### Protein markers of M phenotypes

Stratifying patients into phenotypes using expression-based gene signatures is difficult in a clinical setting. However, measuring circulating proteins in the blood of patients is relatively simpler, and there are standard assays available for this purpose. Thus, identifying protein markers that distinguish these subphenotypes is of great value.

We take advantage of Luminex protein assay profiles from the COMBAT dataset to identify protein markers that are differentially expressed in each M subphenotype and neutrophil subphenotype relative to the others. Unfortunately, we do not have similar data from the other studies we have considered, so the proteins we identify will require further validation before being considered as subphenotype markers.

We provide tables of the differential expression of proteins for each of the M subphenotypes when compared against the “Unc” group as a control. This table is found in the supplementary data file (tab “COMBATLuminexDEPs”). Below we list the proteins that are found to be specifically upregulated in M4 relative to all other categories, and similarly in M2 relative to the others. The first two proteins are more specifically upregulated in M4 (CXCL10, CLEC11a) while the remaining are more specific to M2.

**Table S7:** Luminex proteins that are specific to M subphenotypes.

| protein | M4-Unc | M4-M1 | M4-M2 | M4-M3 | M3-Unc | M2-Unc | M1-Unc | AveExpr | F | adj-PVal |
| --- | --- | --- | --- | --- | --- | --- | --- | --- | --- | --- |
| CXCL10/IP-10/CRG-2 (BR21) (21) high | 2.75 | 2.95 | 1.91 | 3.04 | -0.29 | 0.84 | -0.20 | 12.55 | 15.86 | 2.47E-09 |
| SCGF/CLEC11a (BR76) (76) high | 1.14 | 1.47 | 1.04 | 0.97 | 0.17 | 0.11 | -0.33 | 9.81 | 7.90 | 4.68E-05 |
| IL-6 (BR13) (13) high | 1.09 | 0.01 | -2.13 | -0.21 | 1.30 | 3.22 | 1.08 | 8.25 | 15.15 | 4.10E-09 |
| Lipocalin-2/NGAL (BR21) (21) high | 0.61 | -0.66 | -2.72 | -1.06 | 1.68 | 3.33 | 1.27 | 10.92 | 23.30 | 1.53E-12 |
| CCL20/MIP-3 alpha (BR33) (33) high | 0.48 | -0.01 | -2.29 | -0.44 | 0.93 | 2.77 | 0.49 | 8.52 | 21.94 | 3.45E-12 |

#### Protein Markers of neutrophil phenotypes

In order to identify protein biomarkers of the neutrophil-based phenotypes, we performed differential protein analysis of protein intensities from a Olink panel of 1,472 protein profiles of whole blood from patients with COVID for which whole blood neutrophil data was available (the Massachusetts General hospital cohort: GSE214021). Differential comparisons were made between neutrophil phenotypes using limma. We identified proteins that were significantly differentially expressed across all pairwise comparisons between a given neutrophil phenotype and the other phenotypes. We found that the NeutA phenotype had the most variable expression, and hence although there were proteins that were statistically significant in their differential expression, no protein was overexpressed in this phenotype relative to all of the others with a large (i.e logFC > 0.5) fold change. However, we found 6 proteins that were specifically overexpressed in NeutB (minimum logFC > 0.7) and 15 specific to NeutC. These proteins are listed in Table S8 below. A violin plot of the top 2 proteins for each of these two phenotypes is shown in Figure. S5. Interestingly, CXCL10 is the most significant protein that distinguishes the NeutB hyperinflammatory phenotype, and in parallel also distinguishes the M4 phenotype.

**Table S8:** Pr’from the Olink 1472 panel that are specific to neutrophil phenotypes.

| Assay | NeutB-NeutNormal | NeutB-NeutA | NeutB-NeutC | AveExpr | F | P.Value | FDR |
| --- | --- | --- | --- | --- | --- | --- | --- |
| CXCL10 | 2.40 | 1.65 | 1.53 | 1.28 | 68.36 | 3.93E-39 | 4.37E-36 |
| IFNL1 | 2.03 | 1.30 | 1.60 | 0.91 | 68.01 | 5.94E-39 | 4.37E-36 |
| IFNG | 2.38 | 1.88 | 2.66 | 0.59 | 42.27 | 2.90E-25 | 2.51E-23 |
| CCL8 | 1.30 | 0.96 | 1.14 | -4.03 | 39.48 | 1.03E-23 | 8.00E-22 |
| CXCL11 | 1.19 | 0.95 | 0.99 | 1.07 | 26.03 | 4.92E-16 | 1.96E-14 |
| AGER | 1.06 | 0.86 | 0.72 | -1.29 | 22.78 | 3.98E-14 | 1.01E-12 |

*NeutC.*
| Assay | NeutC-NeutNormal | NeutC-NeutA | NeutC-NeutB | AveExpr | F | P.Value | FDR |
| --- | --- | --- | --- | --- | --- | --- | --- |
| CCL20 | 1.74 | 1.16 | 0.77 | -0.92 | 30.57 | 1.17E-18 | 5.55E-17 |
| CHI3L1 | 1.39 | 1.17 | 0.82 | 0.74 | 21.40 | 2.60E-13 | 6.18E-12 |
| HGF | 1.32 | 0.98 | 1.12 | 0.24 | 21.26 | 3.11E-13 | 7.28E-12 |
| TNFRSF10B | 0.87 | 0.75 | 0.74 | -0.19 | 16.78 | 1.45E-10 | 2.02E-09 |
| REG1B | 1.20 | 1.07 | 0.98 | -0.37 | 16.30 | 2.85E-10 | 3.81E-09 |
| SMOC1 | 1.25 | 0.84 | 1.05 | -0.69 | 14.73 | 2.50E-09 | 2.71E-08 |
| WFDC2 | 0.83 | 0.91 | 0.93 | -0.11 | 13.89 | 7.97E-09 | 7.52E-08 |
| HAVCR1 | 0.86 | 0.76 | 0.75 | -1.38 | 12.89 | 3.21E-08 | 2.67E-07 |
| FGFBP1 | 0.85 | 0.84 | 1.24 | -1.40 | 12.02 | 1.07E-07 | 7.56E-07 |
| CALB1 | 0.87 | 0.84 | 0.85 | -0.66 | 11.81 | 1.45E-07 | 9.98E-07 |
| PTN | 1.30 | 1.03 | 1.66 | 1.10 | 11.38 | 2.65E-07 | 1.71E-06 |
| RSPO1 | 0.76 | 0.73 | 0.83 | -0.63 | 11.02 | 4.36E-07 | 2.65E-06 |
| SFRP1 | 1.56 | 1.23 | 1.80 | -0.12 | 10.92 | 5.00E-07 | 3.01E-06 |
| REG3A | 0.72 | 0.83 | 0.71 | -0.23 | 9.29 | 4.87E-06 | 2.37E-05 |
| MDK | 0.97 | 0.78 | 0.83 | 0.61 | 8.44 | 1.61E-05 | 6.87E-05 |

## Supplementary Figures

**Figure S1:**
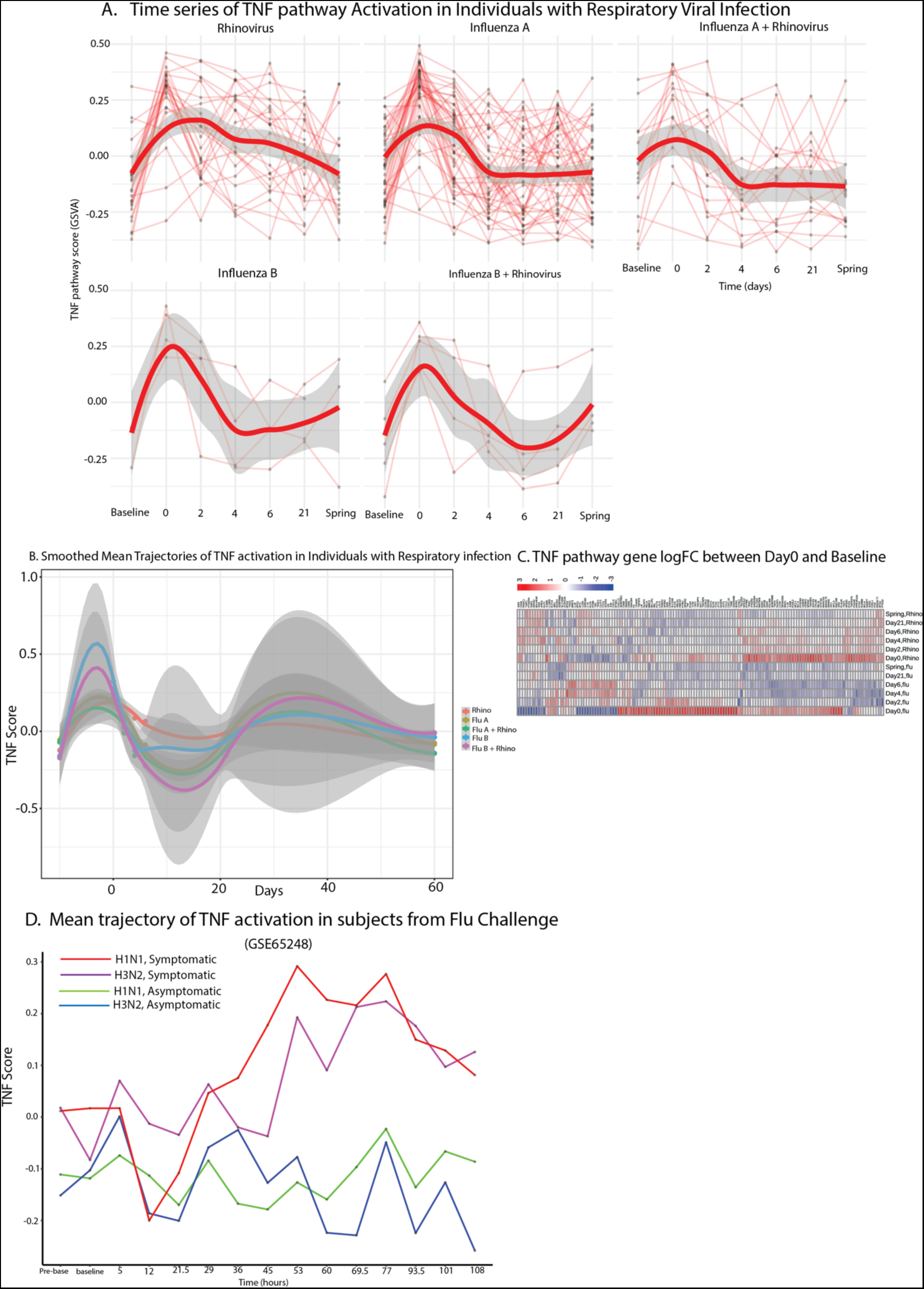
(A) Individual trajectories of TNF activation in patients with respiratory illness from 7 time points. The thick red curve is a smoothed mean value, and the shaded area represents the standard deviation at the timepoints. (B) Smoothed trajectories from (A). (C) Heatmap of TNF pathway gene differential log fold changes comparing day 0 (within 48 hours of acute symptom onset) to a healthy baseline. (D) Is a mean trajectory of TNF activation in subjects symptomatic (n = 20) or asymptomatic (n = 20) upon Influenza challenge followed for 7 days (GSE65248).

**Figure S2:**
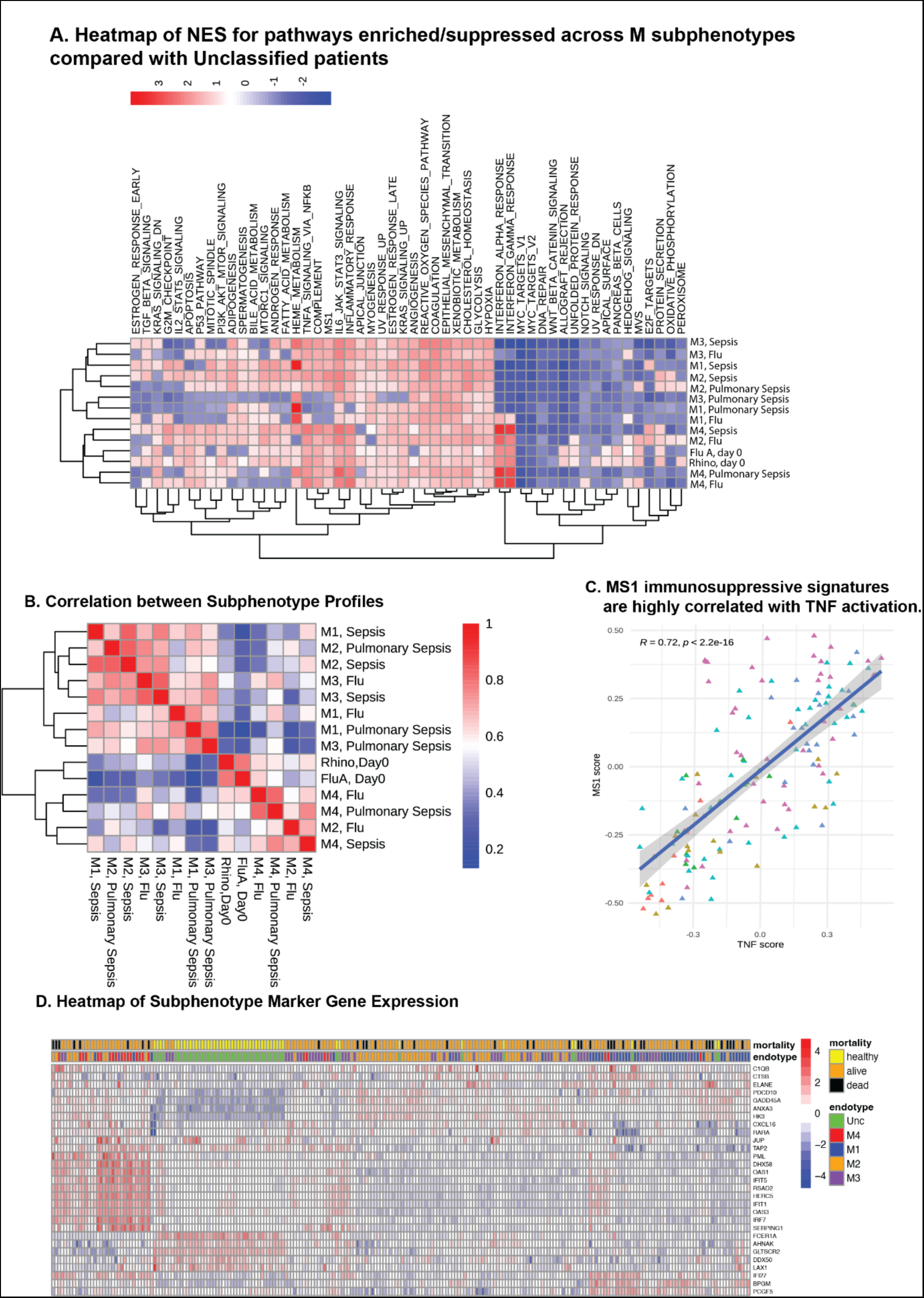
**(A)** Heatmap of NES for the hallmark pathways obtained from a gene set enrichment analysis of all M vs Unc phenotype comparisons. Disease vs Healthy comparisons are also included for completeness. **(B)** Pearson correlations between phenotype NES profiles. The M vs Unc NES profiles cluster together. (**C)** Scatterplot of correlation between TNF signaling and the MS1 MDSC gene signature. (**D)** Heatmap of the expression profiles of M subphenotype marker genes across the pulmonary sepsis dataset (ref). The marker genes do not obviously cluster and correlate with mortality events.

**Figure S3:**
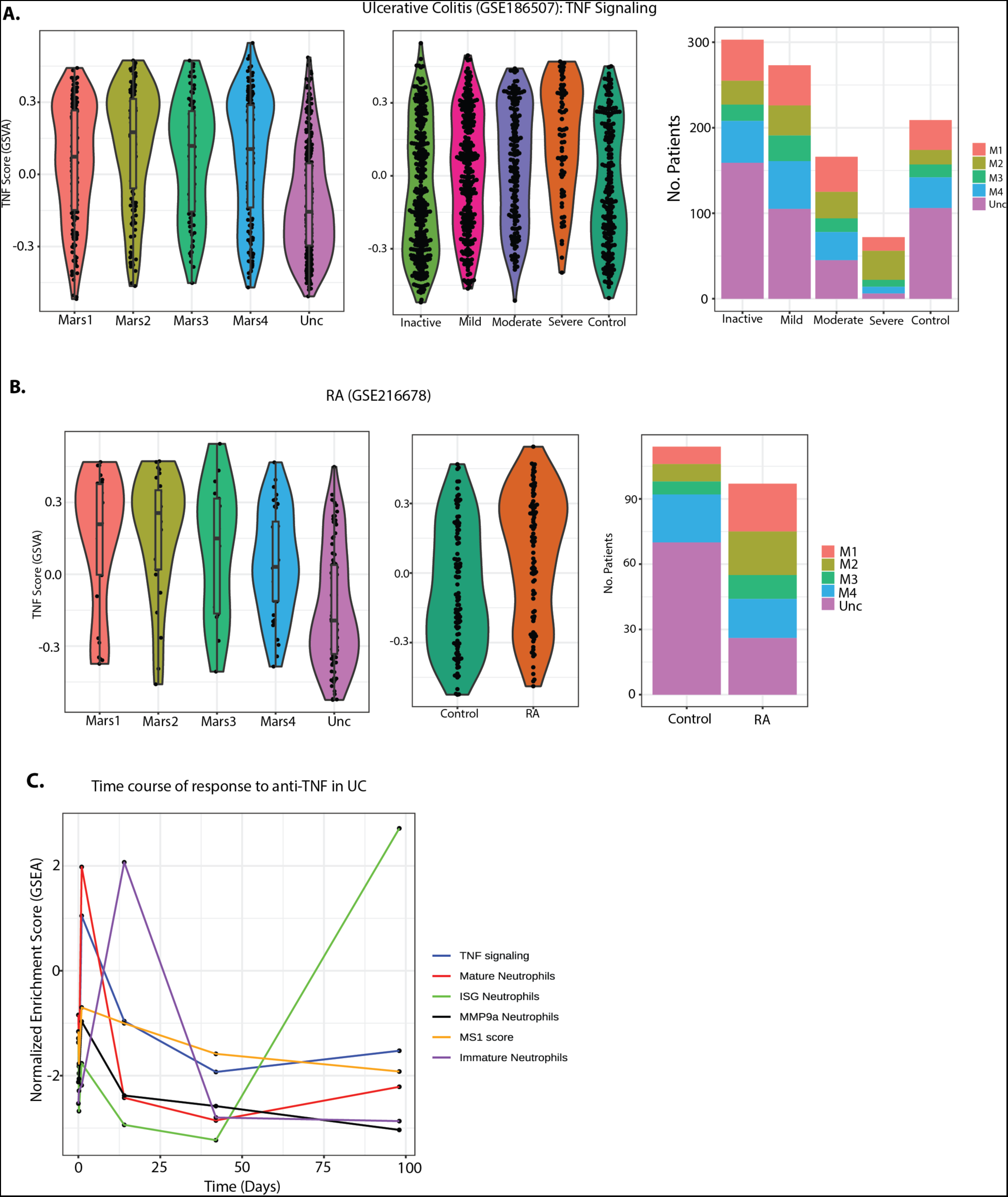
**(A)** TNF pathway activation in UC patients. The M4 subphenotype enriches for patients with high TNF signaling. (**B)** TNF activation in RA patients. RA patients have larger proportions of M1 and M2 than Unc. (**C**) Time series trajectories of GSEA NES scores for the TNF pathway and various neutrophil marker gene sets. The NES values were obtained from gene set enrichment analysis of genes differentially expressed between Responders to anti-TNF therapy compared against non-responders (as defined by the DAS28 score) in patients with Ulcerative Colitis (GSE191328).

**Figure S4:**
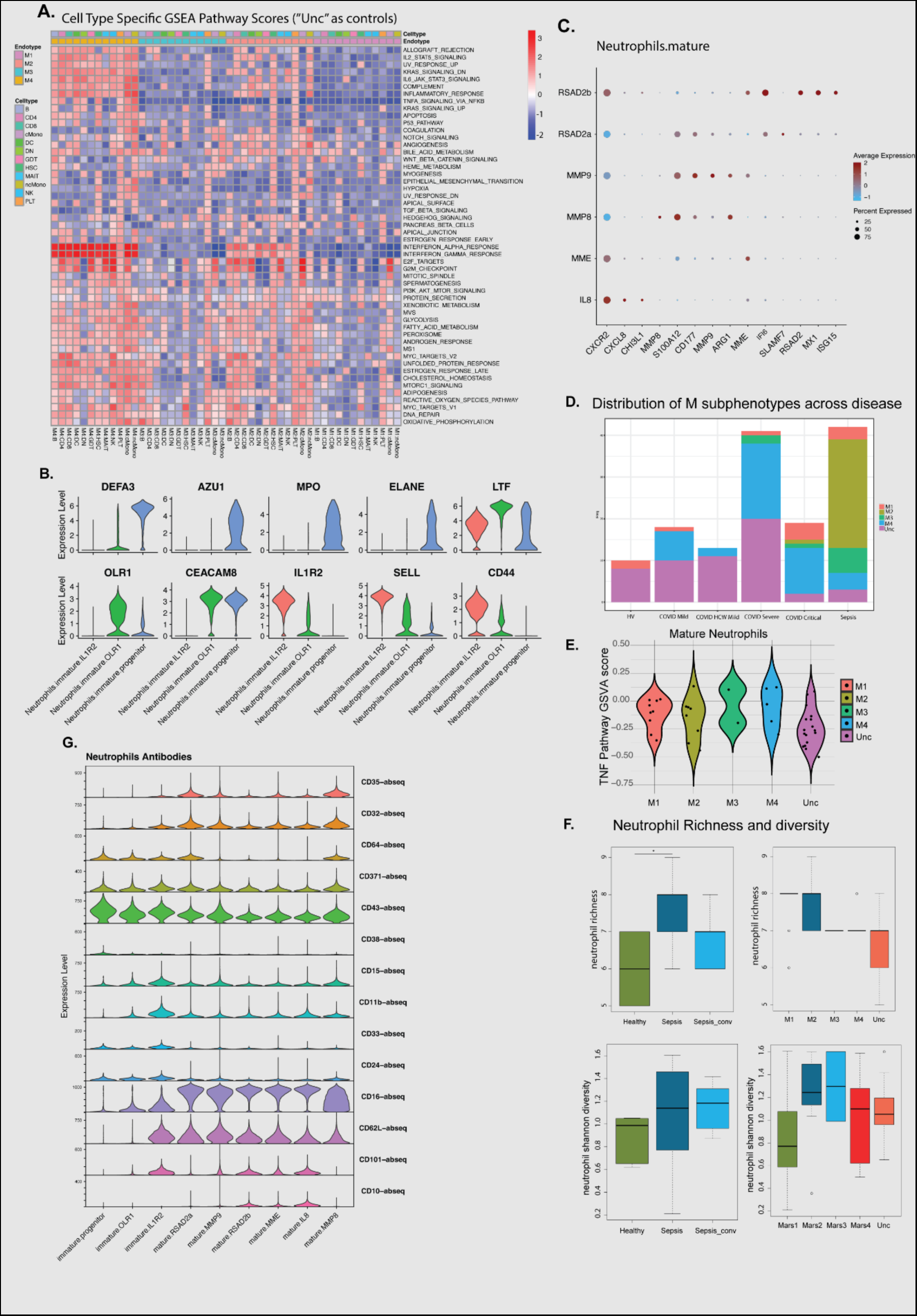
**(A)** Heatmap of Hallmark pathway NES generated from cell type specific differential comparisons of each M subphenotype against the reference “Unc” subphenotype, across the major cell subsets as defined in the COMBAT study. Individual patients from which scRNA profiles were available, were assigned to M subphenotypes based on their whole blood mRNA expression profiles. (**B)** Expression of key neutrophil subset marker genes from the Kwok et. al. scRNA study. (**C**) Percentage of cells expressing and expression magnitude of key marker genes across neutrophil subsets in the Kwok dataset. (**D**) Distribution of M subphenotypes across disease states in the COMBAT COVID dataset. Severe and critical COVID states are enriched for patients with the M4 subphenotype while Sepsis is enriched for patients in the M2 subphenotype. (**E**) TNF GSVA scores across the M subphenotypes when only pseudobulk Neutrophil mRNA expression was considered. The characteristic pattern of TNF activation across subphenotypes is observed. (**F)** Neutrophil Diversity and Richness measured as a function of disease category and M subphenotype for the samples from the Kwok dataset. **(G)** Antibody-derived tags (ADT) expression of surface marker genes from single cell RNA-sequencing across the different neutrophil populations.

**Figure S5:**
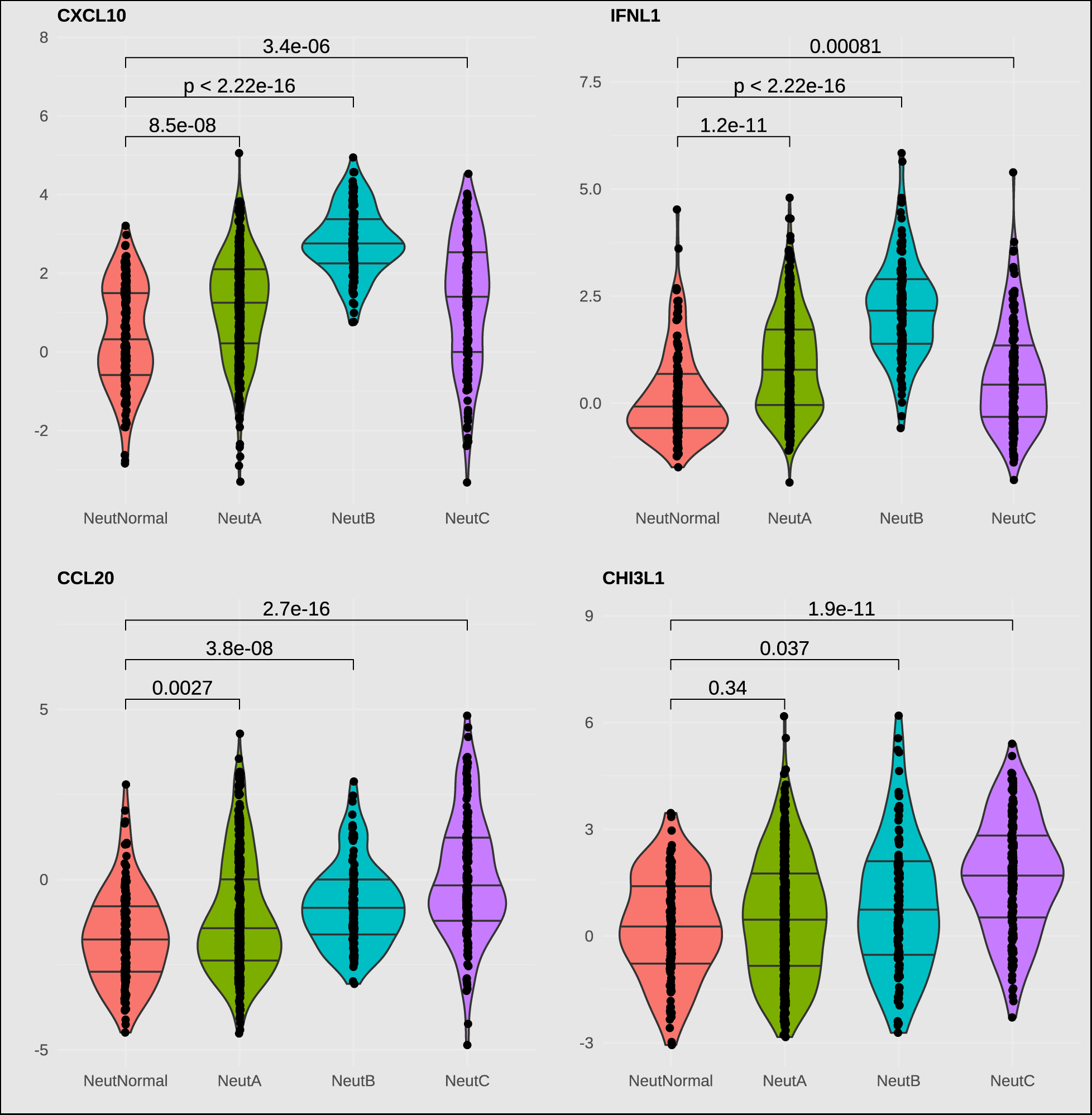
Violin plots of top proteins specifically overexpressed in NeutB (top row) and NeutC (bottom row) relative to the other subphenotypes.

**Figure S6.**
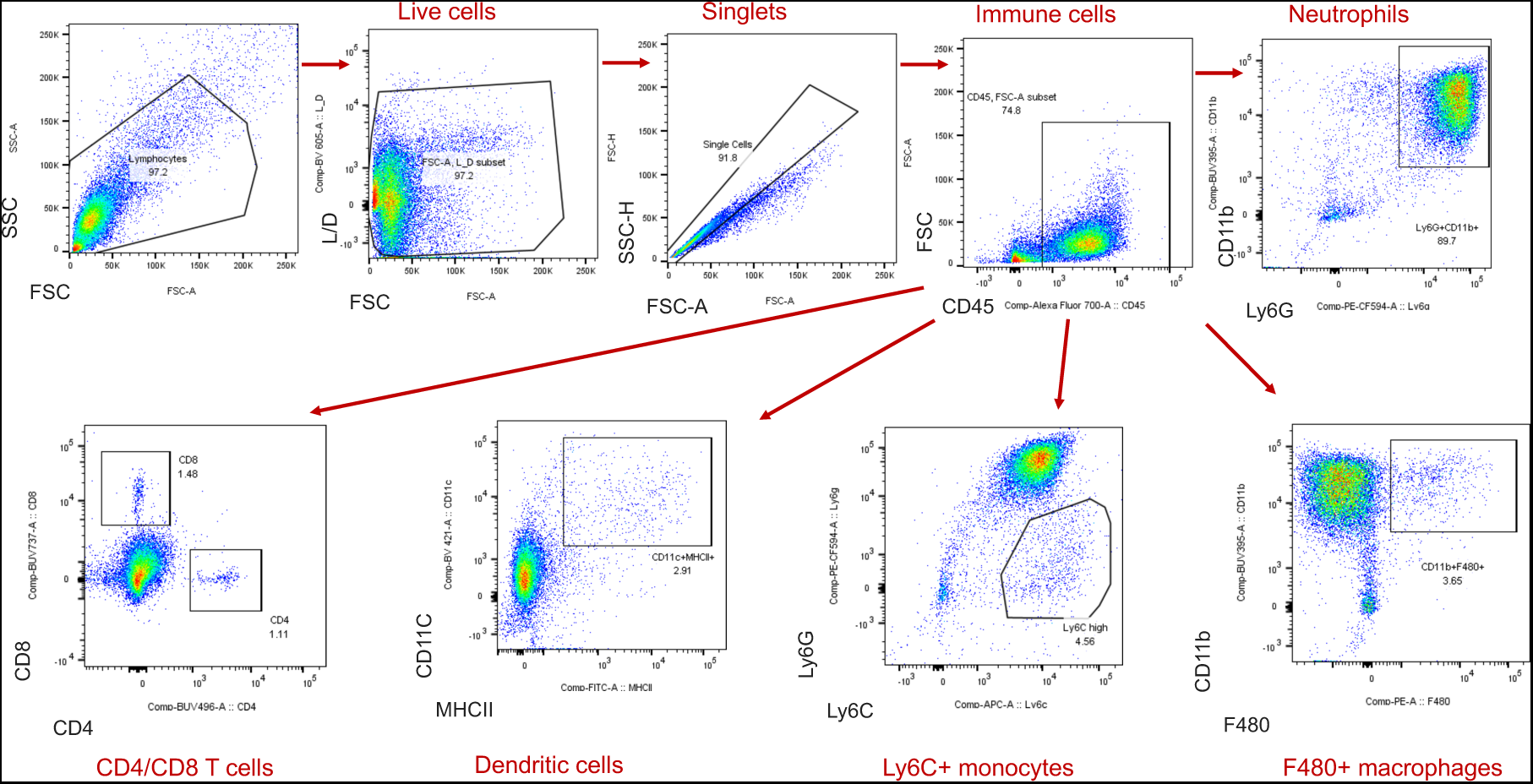
Gating strategy.

## Notes

**Conflict of interest:** All authors were employed by Janssen at the time of the research and may be Johnson and Johnson stockholders.

### Competing Interest Statement

Conflict of interest: All authors were employed by Janssen at the time of the research and may be Johnson and Johnson stockholders.

